# The novel curcumin analogue AKT-100 targets both mutant p53 and STAT3 in gynecologic cancer cells

**DOI:** 10.1101/2025.10.06.680794

**Authors:** Geneva L. Williams, Lane E. Smith, Jamie L. Padilla, Alexander S. Goss, Daisy Belmares-Ortega, Xiangxiang Wu, Jennifer N. Daw, Sumegha Mitra, Prakash Jagtap, Josh K. Monts, Jun-yong Choe, Kimberly K. Leslie

## Abstract

Loss of normal p53 tumor-suppressive activity is common in advanced cancers, where *TP53* mutations produce either p53-null states or highly expressed missense proteins with gain of oncogenic function. Mutant p53 proteins not only lose wild type p53 pro-apoptotic and DNA repair functions but also acquire new activities, including constitutive STAT3 activation. We hypothesize that directly targeting missense mutant p53 instead of downstream pathways is advantageous. Curcumin analogues have been reported to bind mutant p53, motivating the development of AKT-100, a novel analogue designed to restore wild type p53 functionality. AKT-100 binds mutant p53 and STAT3, as shown by fluorescence quenching, and inhibits STAT3 Tyr705 phosphorylation in KLE and COV362 cells. This agent exhibits potent cytotoxicity in gynecological cancer cells with IC_50_ values ranging from 4 nm to 2.51 µM, and demonstrates synergy with the PARP inhibitor olaparib and with chemotherapy by suppressing alternative DNA repair mechanisms associated with therapeutic resistance. RNA sequencing and immunoblotting confirms AKT-100 reactivates wild type p53 transcriptional profiles regulating cell cycle arrest (*CDKN1A*/p21, *GADD45A*), apoptosis (*PMAIP1*/Noxa, *DR5*), and inhibits DNA replication and repair. These findings support AKT-100 as a promising therapeutic agent with the dual capabilities of directly targeting oncogenic mutant p53 and constitutively active STAT3.

**Highlights:**

- AKT-100 is a novel curcumin analogue that functions as a WT p53 reactivator.
- AKT-100 inhibits STAT3 phosphorylation at Tyrosine 705 in endometrial and ovarian cancer cells.
- Cell proliferation is inhibited in multiple ovarian and serous endometrial cancer cell lines with AKT-100 as a single agent.
- AKT-100 is synergistic with PARP inhibitor olaparib and chemotherapy drugs carboplatin and paclitaxel.

**Graphical Abstract:** 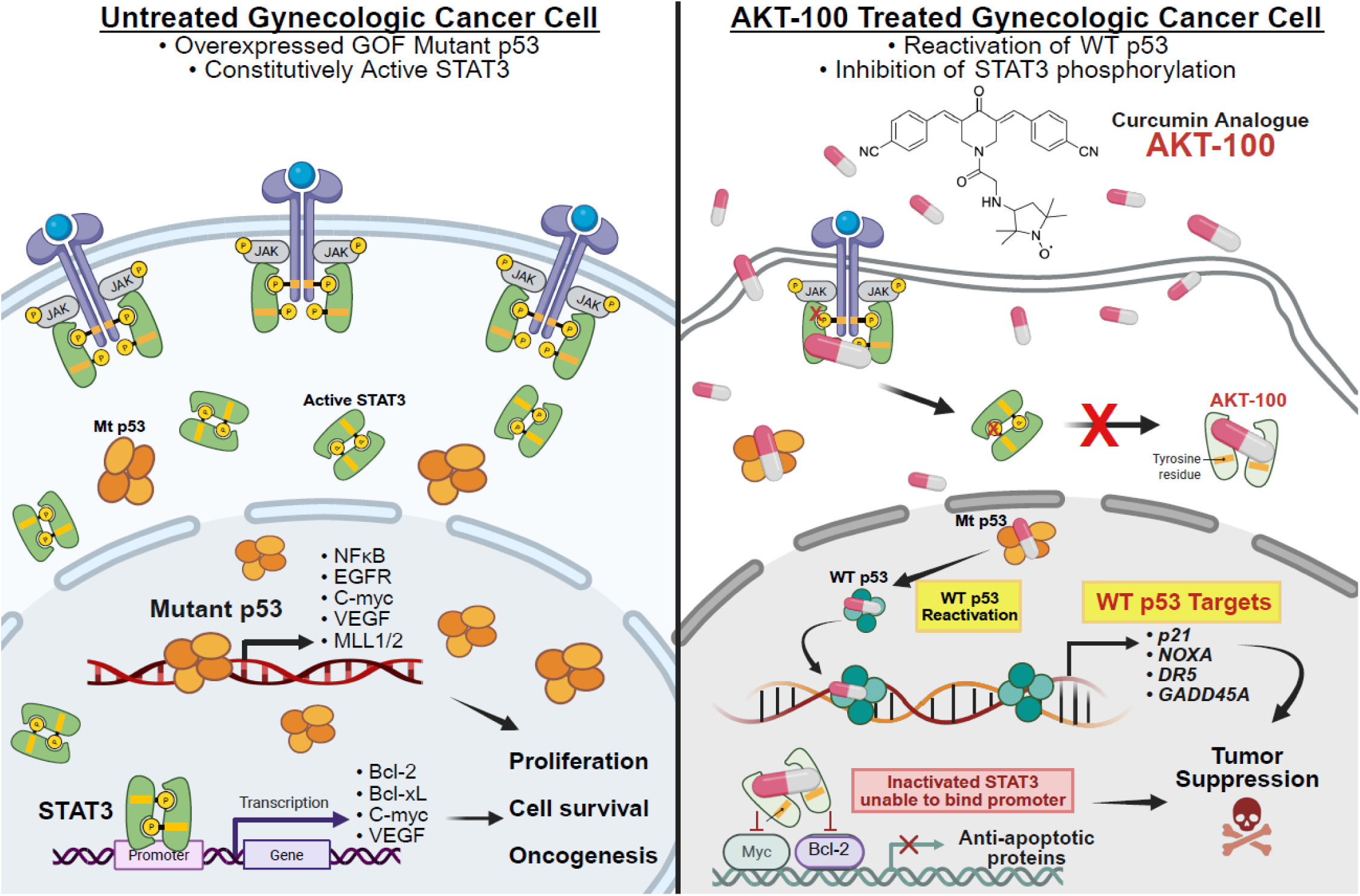

## Introduction

It is well established that mutations in *TP53*, the gene encoding p53, are shared by most types of human malignancies, and these mutations enhance cancer cell growth and survival (1). The evolutionary emergence of p53 homologues is associated with multicellularity, when genome maintenance became a specialized task. Human p53 was discovered more than 40 years ago in the Arnold J. Levine laboratory (2). Initially characterized as a tetrameric transcription factor, normal wild type (WT) p53 also modulates the activity of other factors directly through protein/protein interactions and through DNA binding. It binds to DNA at specific response elements and globally to identify DNA breaks and initiate repair—if repair is not possible, apoptosis is induced (3). Expressed at low, tightly controlled levels in normal cells, p53 is often not expressed (null) or conversely, missense mutated and highly over-expressed in cancer (4–6).

STAT3 (Signal Transducer and Activator of Transcription 3) constitutive activation is associated with poor outcomes in multiple cancers (7, 8). STAT3 signaling is associated with mutant forms of p53, leading to tumors that not only express oncogenic forms of p53, but also are driven by STAT3 hyperactivation (9). STAT3 overactivation is a common finding in advanced cancers; STAT3 is particularly responsible for driving oncogenesis via Myeloid cell leukemia 1 (*MCL1*), cyclin D1 (*CCND1*), MYC proto-oncogene bHLH transcription factor (*c-Myc*), vascular endothelial growth factor (*VEGF*), and Bcl-2 anti-apoptotic family members (*Bcl-xL*, *Bcl-2*) (9). These genes activate multiple pathways associated with cell cycle dysregulation leading to abnormal cell proliferation, differentiation, and resistance to apoptosis. Although multiple indirect STAT3 inhibitors of upstream signaling cascades such as JAK1/JAK2 (ruxolitinib) and SRC (dasatinib) kinases have been approved for clinical use, direct STAT3 inhibitors remain in clinical trials (9, 10). Thus, development of a novel direct inhibitor is potentially an important contribution to the field of oncology by overcoming the challenges of drug resistance caused by constitutively active STAT3.

We also propose that identifying agents that bind to and inhibit the oncogenic properties of missense mutated p53 would be a therapeutically efficacious strategy to prevent and to treat many cancers. Mutations in *TP53* comprise some of the very first alterations in the process of oncogenesis, indicating that targeting these mutant proteins could be a major step towards preventing cancer as well as treating resultant tumors. Importantly, recurrent, missense single amino acid alterations in p53 not only cause the loss of wild type tumor suppression, but result in the acquisition of new oncogenic functionality wherein mutant forms of p53 disrupt nearly all of the normal processes of the cell (11–15). Indeed, the “guardian of the genome” morphs into the “guardian of the cancer cell” (16–18).

The most aggressive and lethal gynecologic malignancies, including uterine serous endometrial cancer and high grade serous ovarian cancer, are characterized by mutations in *TP53* and often highly express the missense mutated forms of the protein that are fundamental to oncogenesis and tumor progression (5, 6, 19). Importantly, the identical single amino acid hotspot p53 mutations, comprising about eight unique, recurrent alterations, occur over and over again in all cancers. These constitute a set of oncogenic “gain of function” proteins that are potentially targetable with p53 reactivator strategies. However, to date, effective p53 reactivator therapies have yet to reach widespread clinical practice. The lack of effective therapeutic options for such p53-mutant cancers constitutes a major gap in medical knowledge and the clinical armamentarium (20).

Gain of oncogenic function mutant p53 oncoproteins significantly enhance angiogenesis (21), inhibit T-cell tumor recognition and permit the over-expression of oncogenes such as *Myc* (12, 22, 23). Based upon preclinical data and with in-depth analyses of clinical studies including GOG 86P (5, 6), NRG GY018 (24) and RUBY (25), mutant p53 has been confirmed as a marker for sensitivity to bevacizumab (anti-VEGFA), pembrolizumab (anti-PD-1) and dostarlimab (anti-PD-1) with significant improvements in both progression free survival (PFS) and overall survival (OS) when added to chemotherapy compared to chemotherapy alone in advanced endometrial cancers. Yet in addition to adding downstream mutant p53-mediated pathway blocking targeted agents to chemotherapy, another therapeutic opportunity would be to target mutant p53 directly, re-establishing the wild type pro-apoptotic functionality as a means to inhibit cancer cell proliferation. Curcumin analogues such as HO-3867 are reported to bind to mutant forms of p53 as well as to STAT3, another important oncogenic driver when over-active, and to inhibit cancer cell proliferation (26–28). Herein, we synthesized a new curcumin analogue, AKT-100, with structural similarities to HO-3867 as well as a series of additional derivatives, AKT-109, 111, 115 and 121. We sought to identify one or more molecules that bind to p53 and reinstate its wild type tumor suppressive activities.

Curcumin itself is widely used as a spice in India and other near Eastern countries which report some of the lowest bowel cancer incidences in the world. It is well known as an anti-inflammatory, anti-angiogenic and anti-oxidant molecule with anti-cancer activity (29, 30). While curcumin is generally considered non-toxic and has been extensively used for medicinal purposes; it is not highly soluble and is poorly absorbed systemically from the gut epithelium. Hence there is a need and an opportunity to create and test curcumin analogues with enhanced solubility for future drug development (31–33).

Opportunities to utilize effective curcumin analogues could range from single agent treatment of malignancies, prophylaxis against cancer development in vulnerable populations, and/or enhancing the activity of standard therapies when given in combination. For example, poly (ADP-ribose) polymerase (PARP) inhibitors commonly used in maintenance for ovarian cancer are most effective in tumors with defects in homologous recombination DNA repair. However, resistance develops when homologous recombination is reconstituted by reversion mutations or other means in cells when exposed to therapy (34). Inhibition of homologous recombination and other DNA repair pathways utilized by cancer cells to escape PARP inhibitor sensitivity by a second agent, such as a curcumin analogue, is a potential mechanism to forestall or reverse PARP inhibitor resistance (35, 36). The same mechanism of tumor DNA repair inhibition could also enhance the sensitivity of tumor cells to chemotherapy. Given the myriad of potential anti-cancer mechanisms of action of curcumin and its analogues, we set out to identify novel therapeutic molecules and to assess their actions and activities. Importantly, we identify agents with the dual function of both reactivating wild type functionality by targeting mutant p53 as well as inhibiting the constitutive activity and oncogenic function of STAT3.

## Materials and Methods

### Synthesis of AKT-100

In collaboration with Akkadian Therapeutics (AKTX, Stoneham, MA), we have identified derivatives of the curcumin analogue HO-3867, such as AKT-100, as ligands that bind to p53. AKT-100 is derived from HO-3867 by removing the toxic functional groups such as tertiary and allyl amines (Figure 1A), which induce the human Ether-à-go-go-Related Gene (hERG) channel blocker activity of HO-3867. The absorption, distribution, metabolism, and excretion (ADME) and drug-like properties of HO-3867 were determined using the computational tool STARDROP (37). The hERG pIC_50_ value of HO-3867 is 6.94 (37). An ideal hERG pIC_50_ value should be <5 for consideration as a safer drug candidate. The proxylamino group on AKT-100 would be expected to satisfy the main pharmaceutical objectives, conferring aqueous solubility (necessary for oral bioavailability, salt formation and oral formulations), and providing long-lasting effects. To identify a highly effective and bioavailable analogue for eventual clinical use, we have designed a library of AKT-100 derivatives which can be tested to identify one or more optimal compounds. Synthetic pathways for the synthesis of AKT-100, AKT-111, AKT-121, AKT-109 and AKT-115 are displayed in Scheme 1 and 2 (Supplemental Figure 1), and synthetic procedures are described in Supplemental Experimental Section. All solvents, reagents and starting materials (i.e. Compound # 1, 2, 3, 4, 12, 14, 15 and 18) required for the synthesis of AKT compounds, were purchased from Sigma-Aldrich, USA. 2-Fluoro-5-((4-oxo-3,4-dihydrophthalazin-1-yl) methyl) benzoic acid (compound # 8, CAS No.: 763114-26-7) for the synthesis of AKT-121(37) was purchased from Ambeed (Buffalo Grove, IL 60089, USA). Proton nuclear magnetic resonance (^1^H-NMR) spectra were obtained from Varian 300 MHz spectrophotometer. Thin layer chromatography (TLC) was carried out on precoated TLC plates with silica gel 60 F-254 and preparative TLC on precoated Whatman 60A TLC plates. ^1^H-NMR of the paramagnetic nitroxide compounds were significantly shifted, and spectral lines were broadened due to the unpaired electron’s magnetic moment. The final compounds were analyzed by mass spectral (MS) analysis data. Flash column chromatography was performed with Merck ultra-pure silica gel (Darmstadt, Germany, 230–400 mesh). Analytical HPLC was performed using a Waters Alliance 2795 series system, equipped with a Waters UV 2996PAD (set at 254 nM) and Micromass MS Quattro LC detector or using Waters Alliance 2690 series system, equipped with the Micromass LCT detector. A YMC-Pack-ODS-AQ (series AQ12505-1546WT, size 150 mm X 4.6 mm, S-5 µM) column was used. A gradient mobile phase starting with 70% water with 0.05% ammonium formate and 30% methanol with 0.05% ammonium formate was employed with a flow rate of 0.8 mL/min. Based on the UV and MS detectors, all final compounds evaluated in the *in vitro* assays showed HPLC purity >95%.

**Figure 1.**
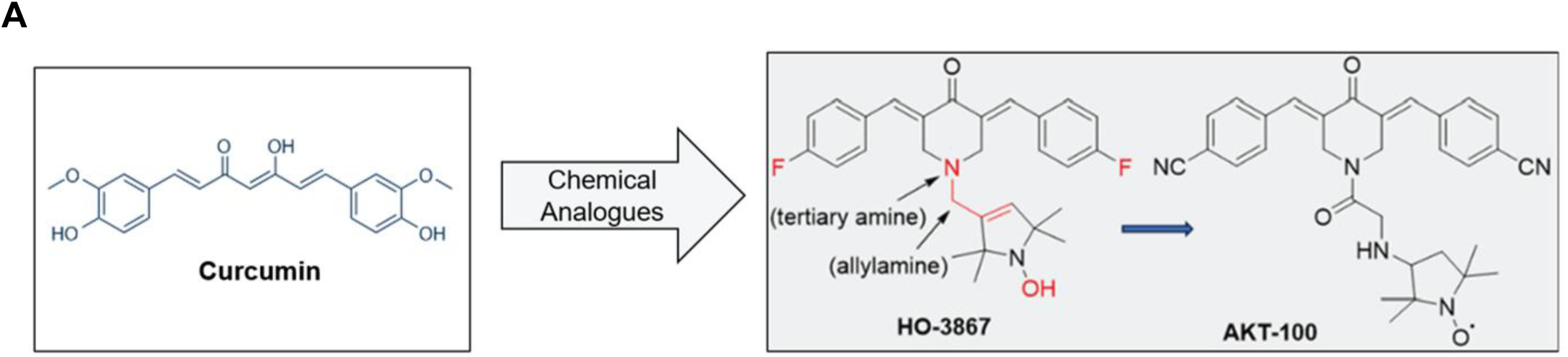
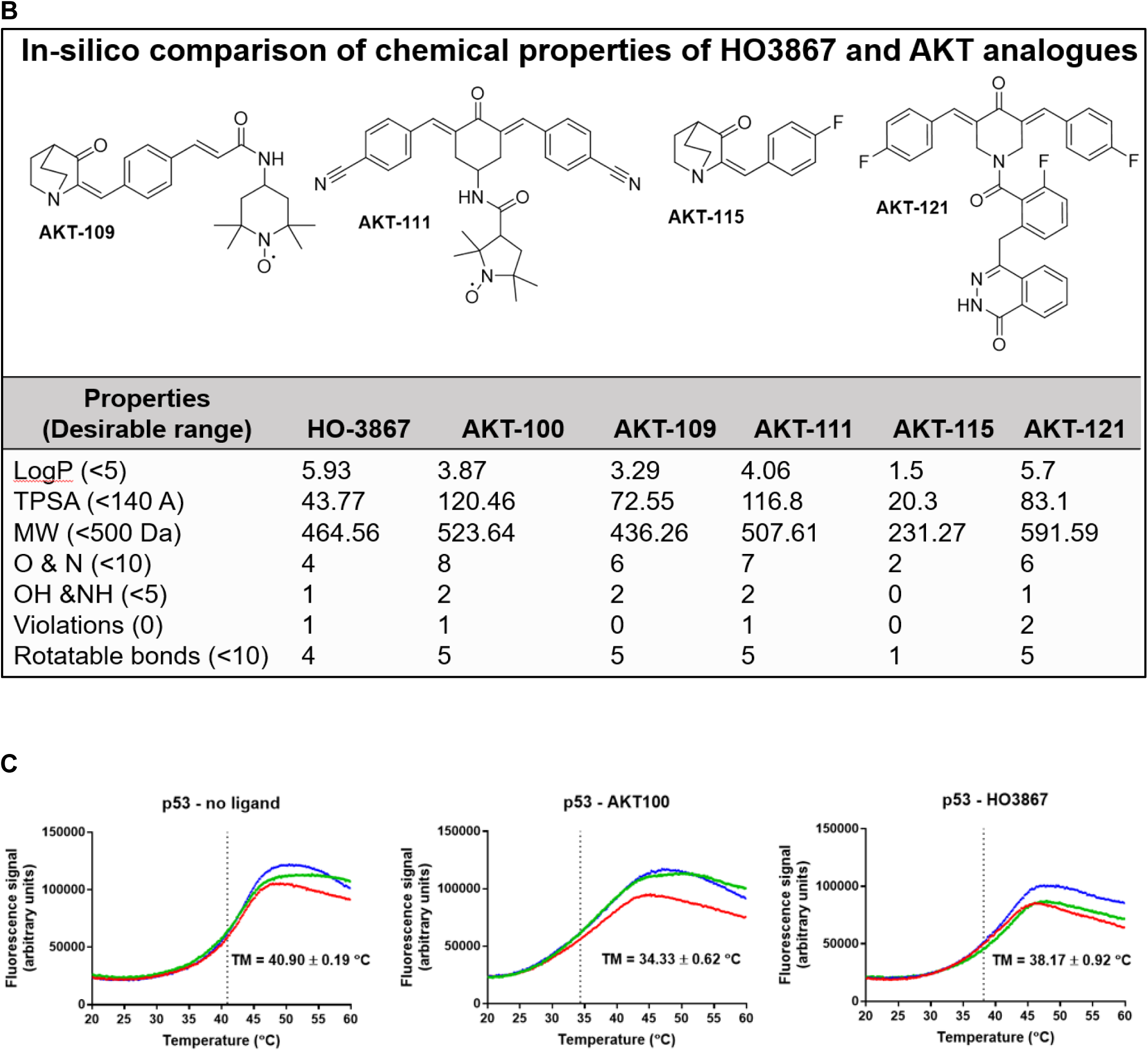
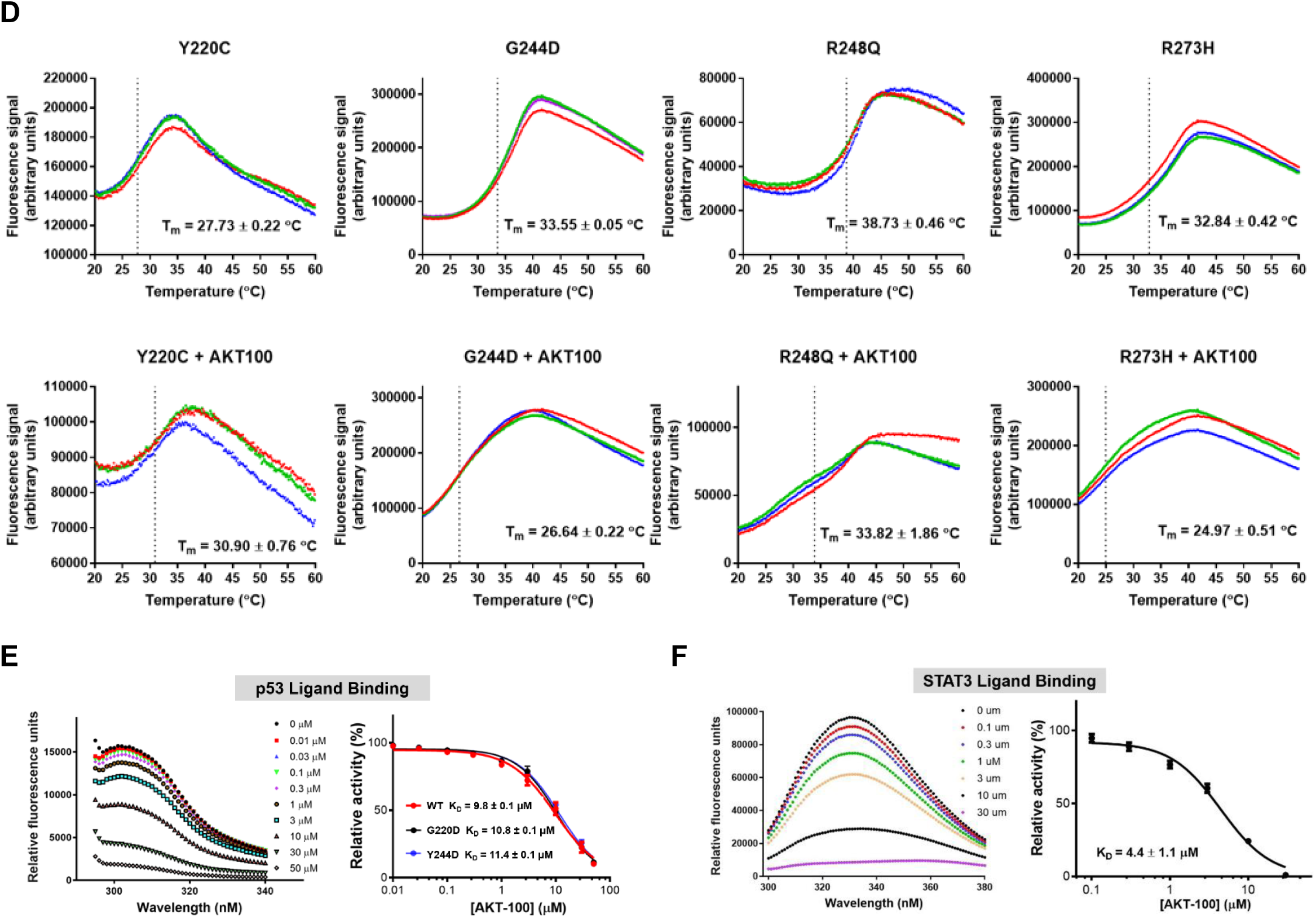
Novel curcumin analogue AKT-100 binds p53 and STAT3 proteins. **(A)** Chemical structure of curcumin and the derivation of novel curcumin analogue AKT-100 from HO-3867. **(B)** Chemical structures of AKT-100 analogues AKT-109, AKT-111, AKT-115 and AKT-121. The chart compares values for all compounds demonstrating favorable *in silico* drug-like properties for solubility and activity compared to HO-3867 and AKT-100. **(C)** Fluorescence changes of SYPRO orange incubated with WT p53 at different temperatures: p53 alone (left), p53 with AKT-100 (middle), p53 with HO3867 (right). **(D)** Fluorescence changes of SYPRO orange incubated with p53 mutants (Y220C, G244D, R248Q, R273H) at different temperatures: p53 mutants alone (top panels), p53 mutants with AKT-100 (lower panels). **(E)** Fluorescence quenching of p53 upon ligand binding. Emission spectra were recorded from 295 nm to 340 nm (excitation at 280 nm) for wild-type p53 in the presence and absence of varying concentrations of AKT-100 (left). Fluorescence quenching of p53 wild type and mutants (G220D and Y244D) at different concentrations of AKT-100 (right). **(F)** Binding of AKT-100 at increasing concentrations to STAT3 was monitored by fluorescence quenching. STAT3 exhibited an intrinsic emission maximum at 330 nm upon excitation at 280 nm (left). Addition of AKT-100 resulted in a concentration-dependent decrease in fluorescence intensity, yielding an apparent dissociation constant (K_D_) of 4.4 ± 1.1 µM. Data represent four independent measurements collected using spectra similar to those shown in panel E.

### Protein purification

The plasmid encoding the human p53 core domain (residues 94–312; AddGene plasmid #24866; (38)) was used as the template. Site-directed mutagenesis was performed using the QuikChange protocol (39) and the resulting constructs were sequence-verified at the University of Chicago Sanger Sequencing Core. The verified plasmid was transformed into *E. coli* C41 cells (40), which were cultured in LB medium. From 10 L of *E. coli* culture, ∼50 mg of p53 protein with ∼80% purity was typically obtained. Purification was carried out using a single step with a Talon (Takara, Kyoto, Japan) affinity column. When the initial purity was below 80%, an additional size-exclusion chromatography step on a Superdex 200 column (Cytiva, Marlborough, MA) was performed. Protein purity was quantified using ImageJ.

### Isothermal Titration Calorimetry (ITC)

ITC was performed on a Nano ITC instrument (TA Instruments, New Castle, DE) at 25 °C. Purified p53 core domain (residues 94–312), and its variants were extensively dialyzed against ITC buffer (50 mM sodium phosphate, pH 7.4, 150 mM NaCl, 1 mM DTT) and degassed immediately before use. The sample cell (300 µL) was loaded with 10 µM p53 protein, while the syringe (50 µL) was filled with 150 µM AKT-100. Each titration consisted of 25 injections of 2 µL with 200 second intervals, under continuous stirring at 300 rpm. Thermograms were integrated, baseline-corrected, and analyzed using NanoAnalyze software (TA Instruments). Binding parameters, including stoichiometry (n), equilibrium dissociation constant (Kd), enthalpy change (ΔH), and entropy contribution (ΔS), were obtained by fitting the data to a multiple site binding model.

### Thermal shift assay (TSA) with SYPRO orange

The fluorescence probe SYPRO orange (λ excitation 490 nm and λ emission 585 nm) binds preferentially to unfolded segments of proteins, and its fluorescence correlates with the degree of protein unfolding (SYPRO orange alone is non-fluorescent). Increasing the sample temperature leads to more protein unfolding which, in turn, is determined as an increase in the fluorescence signal of SYPRO orange. This thermal denaturation assay enables convenient determination of protein melting temperatures using multi-well formats and real-time PCR thermal cyclers (41–43). For each reaction, 5 µg (0.02 mM) of protein was mixed with 10 µM SYPRO Orange in 384-well plates, and fluorescence changes were recorded on a ThermoFisher Studio 6 real-time PCR system. The melting temperature (T_m_) was determined by fitting the fluorescence-versus-temperature curve to a Boltzmann sigmoidal function in GraphPad Prism.

### Fluorescence quenching

Ligand binding perturbs the local environment of tyrosine residues, resulting in decreased intrinsic fluorescence (quenching). Fluorescence changes were monitored as a function of ligand concentration to generate a binding isotherm and determine the dissociation constant (Kd). The experiments shown in Supplemental Figure 2C and D were performed on a Shimadzu RF6000 spectrofluorometer using a single cuvette with excitation at 280 nm. The protein concentration was 0.5 µM.

### Cell Culture

KLE (ATCC # CRL-1622) and OVCAR3 (Cat# HTB-161) cells were obtained from ATCC and grown in RPMI 1640 medium (Gibco Cat# 11875-085 or Corning Cat# 10-040-CV) containing 20% FBS (Gibco Ref# A5670701 or Sigma-Aldrich Cat# 12306C-500ML), 1% penicillin-streptomycin (Gibco Ref# 15140-122), and 1% MEM NEAA (Gibco, Ref# 11140-050). COV362 cells (Sigma Aldrich Cat# 07,071,910) were maintained with DMEM (Gibco Ref# 11965-084) containing 10% FBS and 2 mM glutamine (Gibco Cat# 25030-081). Ishikawa, ECC-1, Hec50co cells (gifts from Dr. Erlio Gurpide, New York University), PEO1 (obtained from Dr. Elizabeth Swisher, University of Washington) and PEO4 (Sigma Aldrich Cat # 10,032,309) cells were grown in DMEM containing 10% FBS and 1% penicillin-streptomycin. AN3CA (ATCC #HTB-111) cells were grown in MEM (Gibco Ref# 11095-080) containing 10% FBS and 1% penicillin-streptomycin. As described by Smith *et al.*, all cells were validated and tested periodically for *Mycoplasma* contamination (44). All cells were grown and maintained at 37°C, 5% CO_2_ and used for fewer than 10 passages for experiments.

### CyQuant Cell Proliferation Assay

Assays were performed as previously described by Smith *et al.* (44) with increasing concentrations of HO-3867 (Selleckchem Cat. #S7501, CAS No. 1172133-28-6), AKT-100 or derivatives AKT-109, AKT-111, AKT-115, and AKT-121 (obtained from AKTX) in fresh media and 100 µL added to triplicate wells. Assays requiring olaparib (Cayman Chemical #10621, CAS No. 763113-22-0) were first drugged with increasing concentrations to determine the IC_50_ in OVCAR3 and COV362 cells. Combined drugging with AKT-100 was then completed using previously obtained AKT-100 IC_50_ values with increasing concentrations of olaparib. Reverse combination assays held olaparib constant at its determined IC_50_ while increasing AKT-100. Proliferation assays using paclitaxel (Cayman Chemical #10461, CAS No. 33069-62-4) were drugged with increasing concentrations to determine the IC_50_ in KLE cells. Combination drugging with AKT-100 was done by maintaining AKT-100 constant at its IC_50_ concentration while increasing paclitaxel. Reverse combination drugging held paclitaxel constant at the previously determined IC_50_ in the presence of increasing concentrations of AKT-100. For experiments using carboplatin (Cayman Chemical #13112, CAS No. 41575-94-4), cells were incubated with increasing concentrations to obtain the IC_50_ value in KLE cells. Combination drugging held one drug constant at the determined IC_50_ value while increasing the other. Graphpad Prism software was used to obtain all IC_50_ values from fluorescence (excitation 485 nm, emission 530 nm) readouts and combinatorial indices were calculated where relevant for combination drugging. All drug treatments were performed in triplicate with two or three independently repeated experiments to confirm results.

### Combinatorial Index Calculations

Upon generating IC_50_ values for single drugs and combination drugging as described above, the below was used to determine synergy.

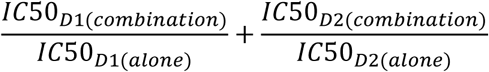

where D1 = drug 1 and D2 = drug 2. Values less than 1 denote synergy.

### RNA Sequencing and Analysis

Cell seeding, RNA collection, sequencing and analysis were completed as previously described by Smith *et al.* (44). Each cell line (Hec50co, Ishikawa, KLE, AN3CA, COV362 and OVCAR3) was treated with AKT-100 at its approximate IC_50_ value (0.50, 0.50, 2.0, 0.10, 0.50, 1.0 µM, respectively) for 24 hours prior to RNA extraction. All samples were processed by the Analytical and Translational Genomics Core at the University of New Mexico. WEB-based GEne SeT AnaLysis Toolkit (WebGestalt) software was used to identify significantly altered pathways using differentially expressed genes. Graphpad Prism software was used to assemble heatmaps displayed in Figure 4B, D and E using Log_2_ fold change values. Additionally, KLE, COV362 and OVCAR3 cells were treated with HO-3867 at their approximate IC_50_ values (4.0, 3.0, and 3.0 µM, respectively) for 24 hours prior to RNA extraction. All RNA sequencing experiments had three biological replicates and differential gene expression analyses were performed by comparing AKT-100 dissolved in DMSO to the DMSO-only control.

### Western blot analysis

KLE and COV362 cells were plated at a density of 800,000 cells/dish, allowed to attach overnight and drugged at 0.1% DMSO, 2 µM AKT-100 or 4 µM HO-3867 for 24 hours. Cells were harvested and lysed for 30 minutes on ice with Pierce RIPA buffer (Thermo Fisher Scientific, Cat# 89900) supplemented with protease inhibitor (Thermo Fisher Scientific Cat# 1860932), followed by centrifugation at 17,000 xg for 10 minutes. BCA protein assays (Thermo Fisher Scientific Cat# 23228, 1859078) were completed in 96-well plates, incubated for 30 minutes at 37°C and absorbance read at 562 nm on a BioTek Synergy Neo2 Plate Reader. 30 µg protein was loaded per well, separated in 4-20% gels (Bio-Rad Cat# 4561094), transferred to PVDF membranes (BioRad Cat# 162-0177) and probed with primary antibody in 3% BSA overnight. Immunoblots were washed three times with 1x TBST prior to addition of horseradish peroxidase (HRP)-conjugated secondary antibody at a 1:1000 dilution in 5% milk for 1 hour. Membranes were washed three times with 1x TBST and imaged after a 1-minute incubation with Pierce ECL Western blotting substrate (Thermo Fisher Scientific Cat# 32106) on a LI-COR Odyssey Fc imager. Antibodies were obtained from Cell Signaling Technology (β-Actin #3700S, p53 (7F5) #2527S, GAPDH-HRP #8884S, PERK #3192S, STAT3 (79D7) #4904S, STAT3 (D3Z2G) #12640S, Phospho-Stat3 (Tyr705) (D3A7) #9145S, Phospho-Stat3 (Ser727) (D4X3C) #34911T, Anti-mouse IgG HRP-linked secondary antibody #7076S, Anti-rabbit IgG HRP-linked secondary antibody #7074S), Santa Cruz Biotechnology (p53 (DO-1) #sc-126, p53 (PAb240) #sc-099, NOXA #sc-56169), EMD Millipore Sigma (Anti-p53 (wild type) clone PAb1620 #MABE339) and Proteintech (p21 #10355-1-AP).

### Immunoprecipitation (IP)

KLE and COV362 cells were plated at a density of 2 million/dish, allowed to attach overnight and drugged at 0.1% DMSO or 2 µM AKT-100 for 24 hours. Cells were harvested and incubated with Pierce IP Lysis Buffer (Thermo Fisher Scientific Cat# 87788) containing Pierce Protease Inhibitor Mini Tablets (Thermo Fisher Scientific Cat# A32953), followed by BCA protein assay. 500 µg lysate was incubated overnight with 2 µg PAb240 or PAb1620 antibody then added to 20 µL Pierce Protein G Magnetic Beads (Thermo Fisher Scientific Cat# 88847) for two hours. Magnetic beads were washed three times with 1xPBS containing protease inhibitor. Proteins were eluted with 20µL of 2x Laemmli Sample Buffer (Bio-Rad Cat# 1610737) and boiling at 100°C for 10 minutes then analyzed by Western blotting with p53 (DO-1) as mentioned above.

### Cell Cycle Analysis via Flow Cytometry

Ishikawa and KLE cells were plated at a density of 2 million/dish, allowed to attach overnight. Ishikawa cells were drugged with 0.1% DMSO or 1 µM AKT-100 for 24 hours. KLE cells were drugged with 0.1% DMSO or 2 µM AKT-100 for 24 hours. 1 million cells from each treatment was collected in 1xPBS and pelleted at 3300 RPM for 3 minutes. BioLegend Zombie NIR Fixable Viability Kit (Cat# 423105) protocol was followed with an incubation time of 30 minutes in the cell stain then washed with 1mL of 2% FBS in 1xPBS. Cells were pelleted, wash buffer aspirated and resuspended in 50 µL of 2% FBS in 1xPBS. 1mL of cold, pre-chilled 70% ethanol was added to the cells, incubated overnight at 4°C. Cells were pelleted at 4000 RPM for 3 minutes, washed twice with 2% FBS in 1x PBS, then 500 µL of 1 µg/mL DAPI (4’,6-Diamidino-2-Phenylindole, Dihydrochloride, Invitrogen™ Cat# D1306) in 0.1% Triton X-100 (Sigma-Aldrich, CAS No. 9036-19-5, Cat# T8787-50ML) was added and incubated at room temperature for 30 minutes. Cells were filtered and samples were run on a Beckman Coulter Cytoflex S cytometer and analyzed using BD FlowJo v10.10.0 software. Based on DAPI-staining of DNA content, G1, S, and G2 phases were reported using Watson Pragmatic modeling, following the exclusion of debris, doublets, and dead cells using a fixable viability dye. Two replicate samples were completed for each cell line. All samples were processed and analyzed by the UNM Flow Cytometry Shared Resource Core.

### Statistical Analyses

IC_50_ values from the cell lines were determined using GraphPad Prism software, version 9.3.1, by applying a nonlinear regression analysis (*[Inhibitor] vs. normalized response* curve) on the normalized fluorescent values. Western blots were quantified using Image J software and entered into GraphPad Prism software followed by analysis with One-way ANOVA.

## Results

### Novel curcumin analogue AKT-100 binds WT and mutant p53 protein

As displayed in Figure 1A and B, AKTX has synthesized the compounds to feature combinations of multiple double bonds, acting as Michael acceptors or interacting with cysteine residues. Our aim in developing these compounds was to maintain their efficacy as well as to address some of the following: (i) increased water solubility; (ii) improved oral drug delivery; and (iii) improved DMPK parameters. AKTX synthesized multiple analogues for our preclinical validation studies, performing pharmacokinetic (PK) and pharmacodynamic (PD) studies to optimize dosing and administration of lead molecules.

We compared the chemical properties and chemical structures of AKT-100 and other AKT derivatives with HO-3867 for their solubility and oral bioavailability (Figure 1B). The essential parameters to predict molecular solubility and cell membrane permeability for optimizing *in vivo* absorption, such as lipophilicity (LogP), topological polar surface area (TPSA) and molecular weight (MW) were within the optimal range. The desired optimal range of the chemical properties are listed in parentheses in Figure 1B (45, 46). TPSA values help determine a molecule’s surface area polarity where higher values are indicative of increased polarity with decreased lipophilicity and overall poor membrane permeability. In addition, a molecule’s optimal molecular weight should be less than 500 g/mol and LogP value less than 5 to achieve desirable oral bioavailability (47). LogP values show that AKT-100 and its derivatives are less than 5 with variable molecular weight’s, indicating potential for increased bioavailability compared to HO-3867. Also displayed in Figure 1B are the chemical structures of AKT-100 derivatives used in our studies to identify the most effective compound that both binds to mutant p53 to reinstate the normal conformation with normal, pro-apoptotic gene transcription and tumor suppression before leading to future *in vivo* experiments.

To validate the lead compound, AKT-100, for binding to the wild type p53 compared to the most common, recurrent mutant forms of p53 (R175H, Y220C, G244D, R248Q, R273H, D281H), we used ITC, the protein melting temperature and protein fluorescence. Purified WT or mutant p53 protein was obtained as described in the methods section and confirmed by SDS PAGE analysis (Supplemental Figure 2A). ITC is a biophysical, label-free analytical technique for determining the binding of molecules in solution (48, 49). The ITC instrument measures the change in the energy of two given interacting molecules as the heat release in the solution, allowing quantitative determination of thermodynamic parameters such as: enthalpy (H), entropy (ΔS), and Gibbs free energy (ΔG), and the equilibrium association constant (Ka), from which the equilibrium dissociation constant Kd can be calculated. ITC data (Supplemental Figure 2B) showed that AKT-100 binds p53 with dissociation constants (Kd) of 5.1 µM and 523 µM, consistent with a multiple sites binding model. The interaction is endothermal, with a predicted stoichiometry of 0.6 and 0.9 (ligand to protein), which reflects sequential ligand binding events and protein dimerization.

Next, the fluorescence probe SYPRO orange was used to determine the protein melting temperature (Tm, denaturation midpoint) as Tm changes upon incubation with a prospective ligand indicate ligand binding (41, 42, 50). The melting temperature of p53 WT and mutants (G244D, R248Q, R273H) decreased upon incubation with AKT-100 or HO-3867 (Figure 1C and D). However, the Tm of mutant p53 variant Y220C increased, possibly indicating greater thermodynamic stability.

Fluorescence quenching is a method to check ligand binding using protein fluorescent residues, such as tryptophan (Trp), tyrosine (Tyr), or phenylalanine (Phe). The changes in the environment of the fluorescent residue induced by ligand binding alters the fluorescence signal (51–54). The core domain of p53 has one Trp, eight Tyr and nine Phe residues. Figure 1E (left panel) shows an emission signal of p53 at 308 nm after sample excitation at 280 nm; the addition of the ligand decreases the emission signal in a concentration-dependent manner. The resulting K_D_ values from the nonlinear regression fit of the data were 9.8 ± 0.1 µM, 10.8 ± 0.1 µM, and 11.4 ± 0.1 µM, for the p53 WT, G220D and Y244D, respectively (Figure 1E, right panel). In Figure 1F we also used this method to show AKT-100 binds to STAT3 with a K_D_ value of 4.4 ± 1.1 µM, indicating higher affinity than p53.

Thus, we have demonstrated binding of AKT-100 to both WT and mutant p53 proteins as well as STAT3, indicating target specificity. Our next step was to demonstrate drug efficacy in cancer cell models and the re-establishment of many functional elements of WT p53, such as cancer cell cycle arrest and induction of the pro-apoptotic transcriptome.

### AKT-100 is more therapeutically effective compared to HO-3867 in multiple cell models

Cell proliferation assays identified IC_50_ values in the low µM - nM range for both AKT-100 and the parent molecule HO-3867 in ovarian and endometrial cancer cell lines harboring p53 mutations. According to Figure 2A and 2B, six cell lines had IC_50_ values ranging from 4-390 nM. Figures 2C-I compare HO-3867 versus AKT-100 treatments in PEO1, ECC-1, KLE, COV362, OVCAR3, Hec50co and Ishikawa cells, with graphs displaying growth inhibition at lower concentrations of AKT-100. These data demonstrate the effectiveness of the novel curcumin analogue AKT-100, however we further asked whether other AKT derivatives are effective at killing cells in our cell models of interest.

**Figure 2.**
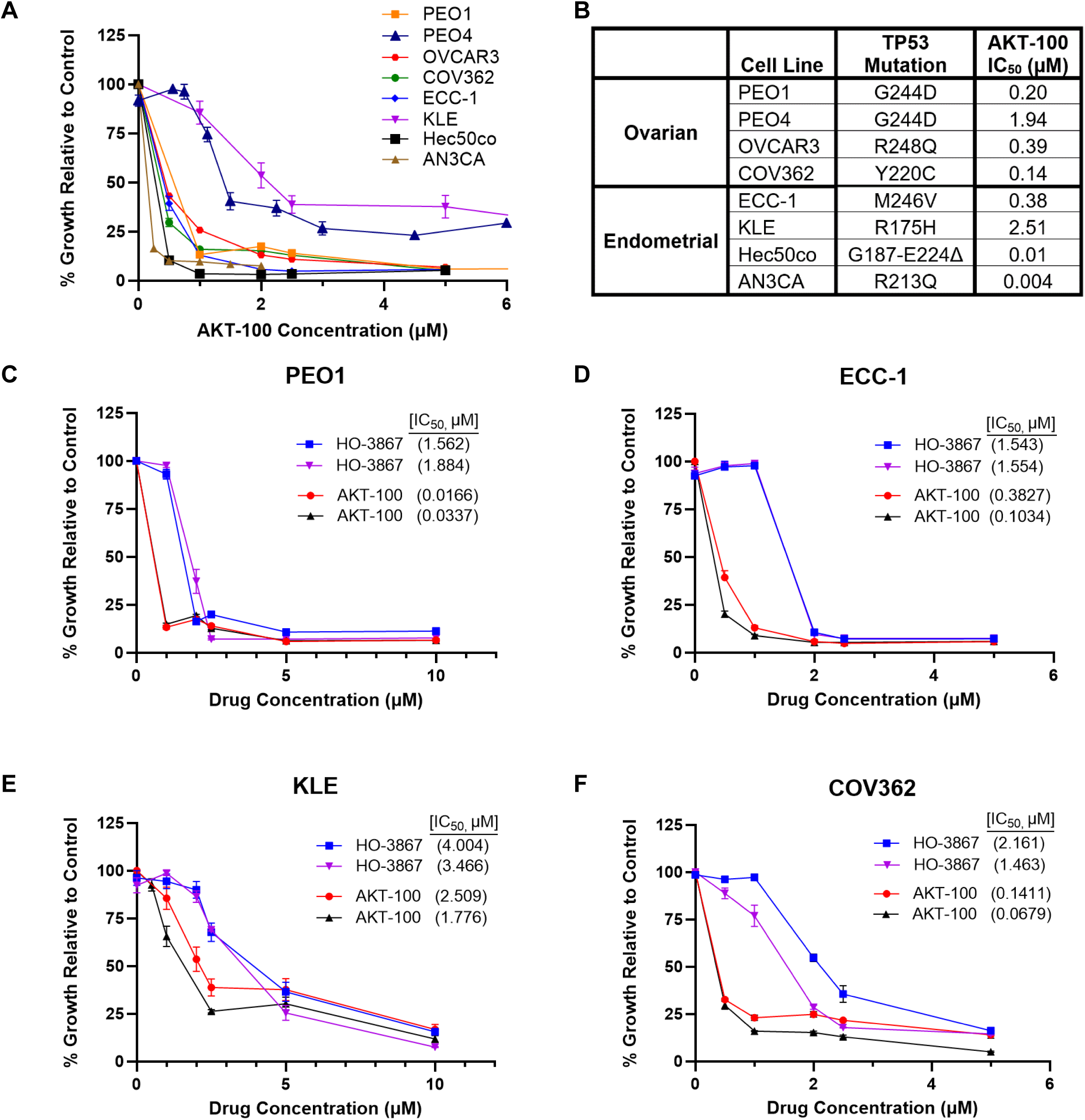
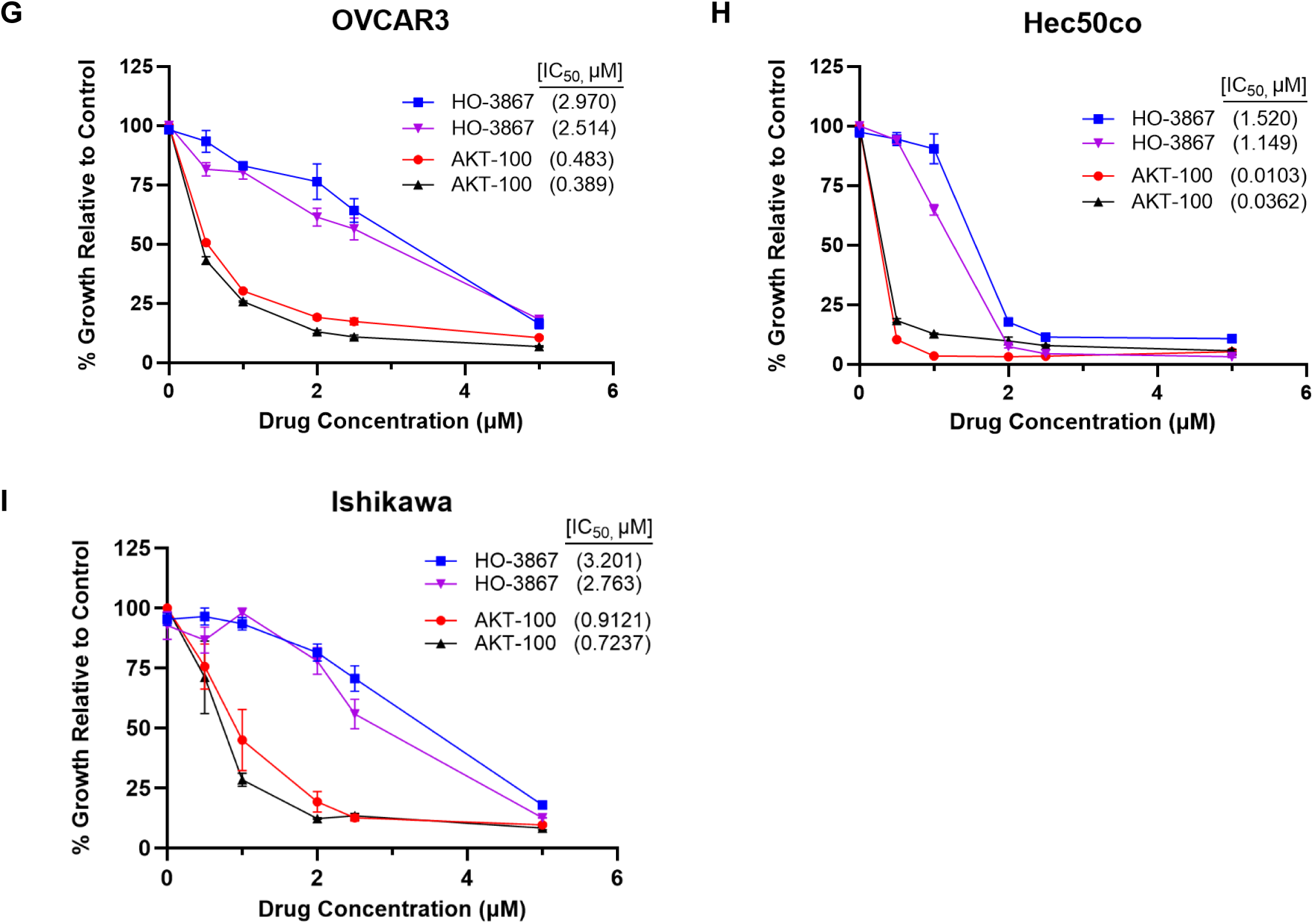
AKT-100 is more effective at killing cancer cells compared to HO-3867. **(A)** CyQUANT cell proliferation assay results of ovarian and endometrial cancer cell lines drugged for 72 hours with increasing concentrations of AKT-100. Error bars represent +/- standard error of the mean (SEM) for triplicate wells. **(B)** Table displaying the IC_50_ values obtained for each cell line treated with AKT-100 and corresponding TP53 amino acid mutations. **(C-I)** Proliferation assays for PE01, ECC-1, KLE, COV362, OVCAR3, Hec50co and Ishikawa cells after drugging with increasing concentrations of HO-3867 versus AKT-100 for 72 hours. Two biological replicates with IC_50_ values in parentheses are shown for each cell line and drug treatment on the same graph.

### AKT-111 and AKT-121 also demonstrate potential activity against cancer cells

Curcumin analogue AKT-111 was more effective at killing KLE, OVCAR3 and PEO1 cells when compared to AKT-109 and AKT-115. The calculated IC_50_ values for AKT-111 remained in the low micromolar range (3.526 µM for KLE, 2.60 µM for OVCAR3, 1.69 nM for PEO1), whereas AKT-109 and AKT-115 failed to reach an IC_50_ in our experiments (Supplemental Figure 3A-C). Upon comparison of AKT-100 with AKT-121, AKT-100 again proved to be more efficient at inhibiting cell proliferation; however, the IC_50_ for AKT-121 remained in the low micromolar range as well for KLE and COV362 cells (Supplemental Figure 3D-E).

### Tumor suppressive p53 transcriptional effects are reinstated after treatment with AKT-100

KLE cells, harboring the oncogenic p53-R175H mutation, were exposed to 2.0 μM AKT-100 for 24 hours, and RNA sequencing revealed the re-enforcement of cell cycle checkpoints by the upregulation of *CDKN1A* (p21) and *GADD45* (GADD45) (Figure 3A). Proliferative signaling downstream of *ATM* (ataxia telangiectasia mutated), an important serine/threonine protein kinase stimulating cell growth, was also downregulated by AKT-100 (Figure 3A). Simultaneously, genes within the apoptotic pathway were induced that predicted for programmed cell death (Figures 3A, 3B). In our *in vitro* models, AKT-100 and HO-3867 restored WT tumor suppressor function in otherwise oncogenic mutant p53 by reducing cancer cell replication and promoting apoptosis. However, AKT-100 proved to be more effective than HO-3867 in OVCAR3 cells at upregulating *CDKN1A* (p21) and *GADD45* transcripts (Supplemental Figure 5). Cell cycle evaluation by flow cytometry confirmed the reinforcement of the G2/M checkpoint in KLE cells with the p53-R175H mutation after AKT-100 treatment, likely linked to the upregulation of gene transcription of *CDKN1A* and *GADD45* (Figures 3B and 3E). In addition, Western blot analysis demonstrated that AKT-100 increases the protein expression of p21, NOXA and PERK in KLE and COV362 cells (Figure 3C) compared to the DMSO control and HO-3867 treated cells. Figure 3B displays an increase in p21 and GADD45 (left panel) in all cell lines harboring *TP53* mutations, while there is also a decrease or no induction of DNA repair genes (middle panel) across these same cell lines upon treatment with AKT-100. The downregulation of the majority of genes involved in multiple DNA repair pathways (middle panel), with the exception of *XRCC4* and *LIG4* central to non-homologous end joining (NHEJ), portends that AKT-100 may be useful to enhance the effects of standard therapies such as PARP inhibitors and chemotherapy, as we test below.

**Figure 3.**
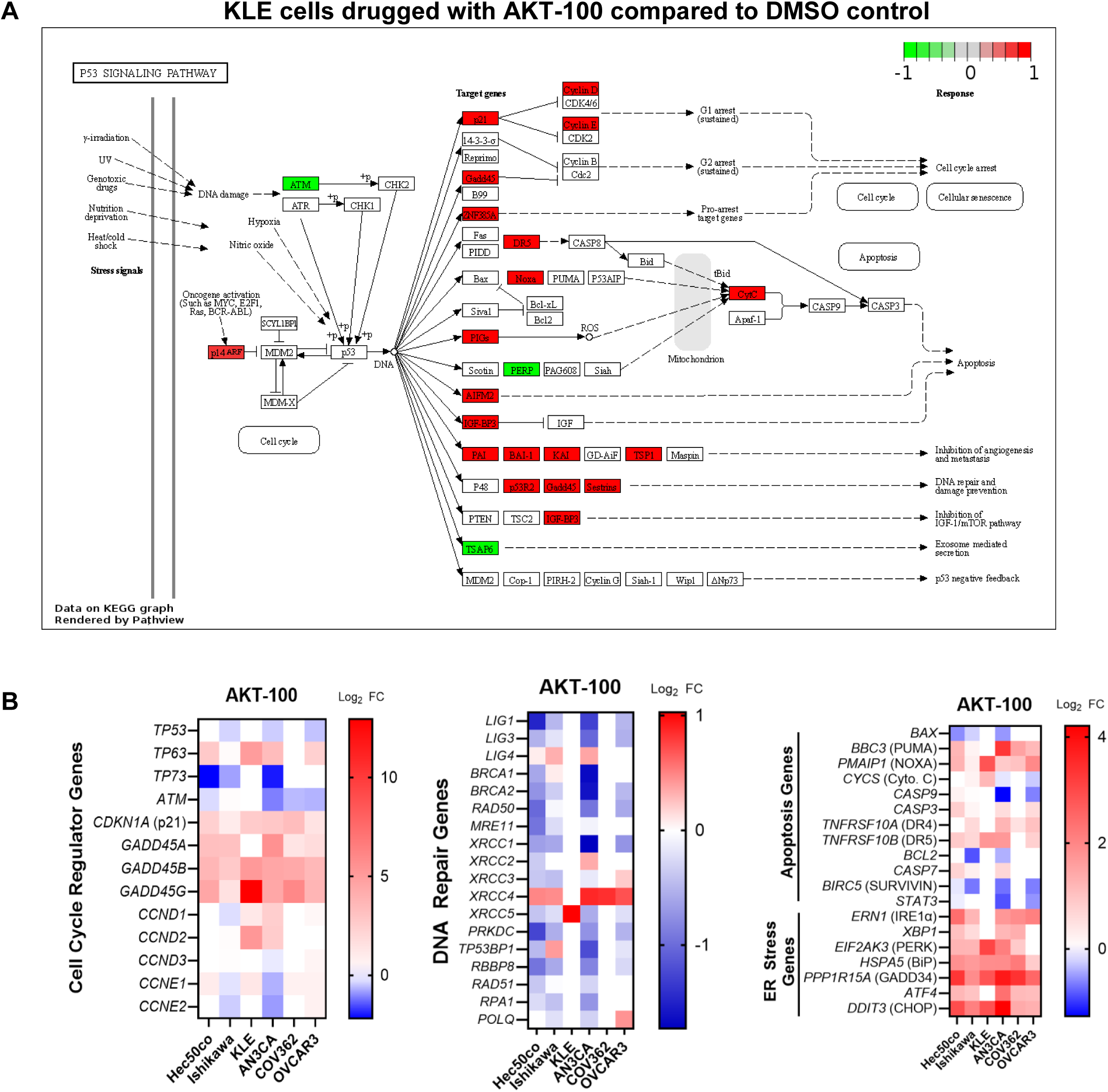
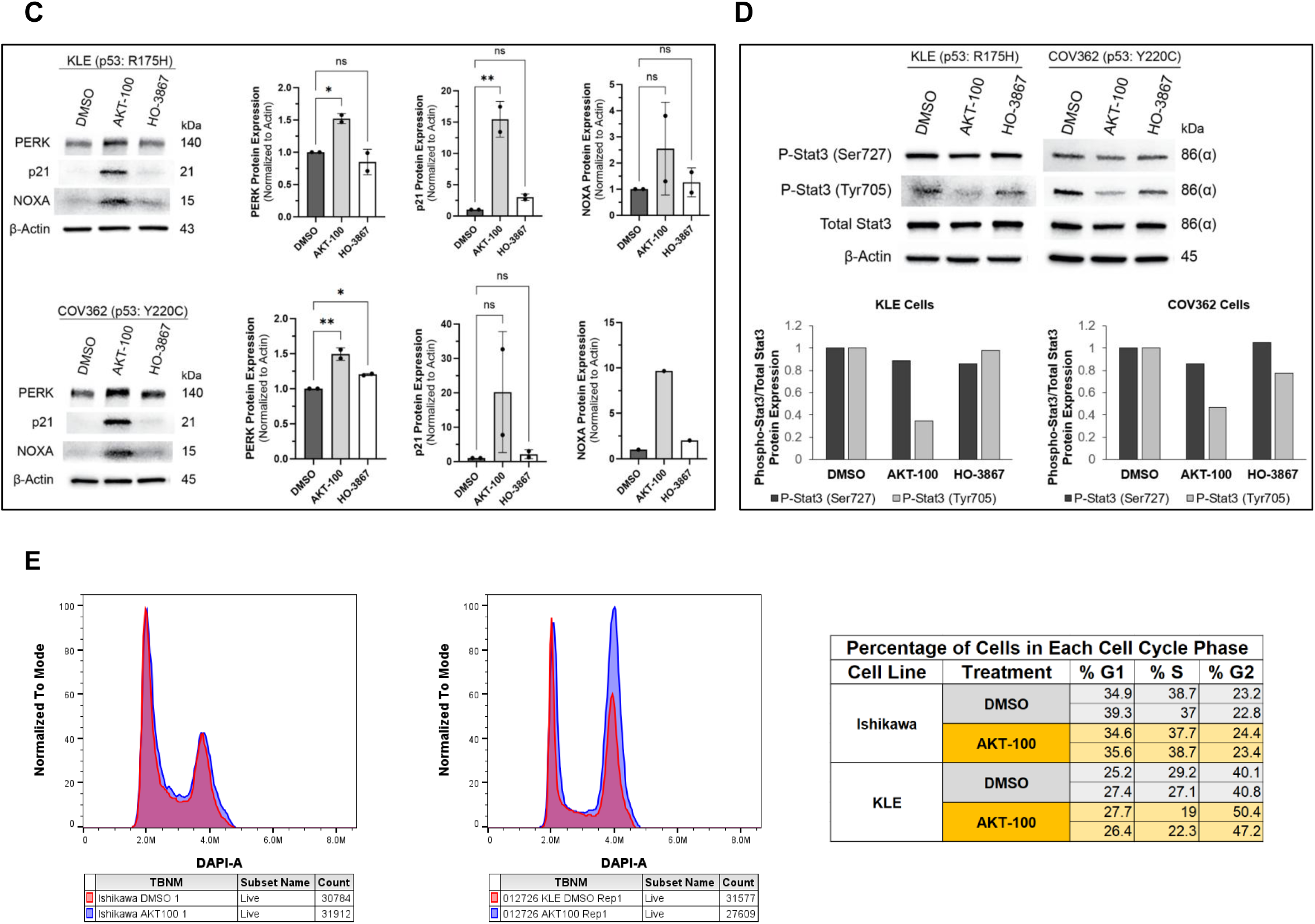
Tumor suppressive p53 transcriptional effects are reinstated after treatment with AKT-100. **(A)** Pathview analysis of RNA sequencing results in KLE cells drugged with AKT-100 showing upregulation of cell cycle regulating and apoptosis genes. Highly upregulated genes are highlighted in red and downregulated genes in green. **(B)** Heatmaps displaying upregulation (red) or downregulation (blue) of cell cycle regulating genes (left panel), DNA repair genes (middle panel) or apoptosis and ER stress genes (right panel) after treatment with AKT-100 in Hec50co, Ishikawa, KLE, AN3CA, COV362 and OVCAR3 cells. Note: For heatmaps, the scale bars represent log_2_ fold change values of transcripts in AKT-100 treated cells compared to DMSO control treated cells. **(C)** Western blot analysis confirming upregulated protein expression of p21, PERK, and NOXA in KLE and COV362 cells treated with AKT-100 for 24 hours. One-way ANOVA was performed to compare treatments to DMSO, where statistical significance is indicated as follows: *, p < 0.05; **, p < 0.01; ns, not significant. **(D)** Western blot showing downregulation of phosphorylated Stat3 with AKT-100 treatment only at Tyrosine 705 in KLE and COV362 cells. **(E)** Ishikawa (left panel) versus KLE (middle panel) cell cycle analysis using DAPI staining of DNA content after 24-hour treatment with AKT-100. Ishikawa cells were treated with 1 µM of AKT-100 and shows similar profiles. KLE cells had a dosage of 2 µM AKT-100 and shows an increase of cells in G2 arrest. (Cells were drugged at the previously determined IC_50_ values.) The table compares the percentage of cells in each cell cycle phase (G1, S or G2) in Ishikawa versus KLE cells upon treatment with AKT-100. Two replicates were completed for analysis by flow cytometry.

**Figure 4.**
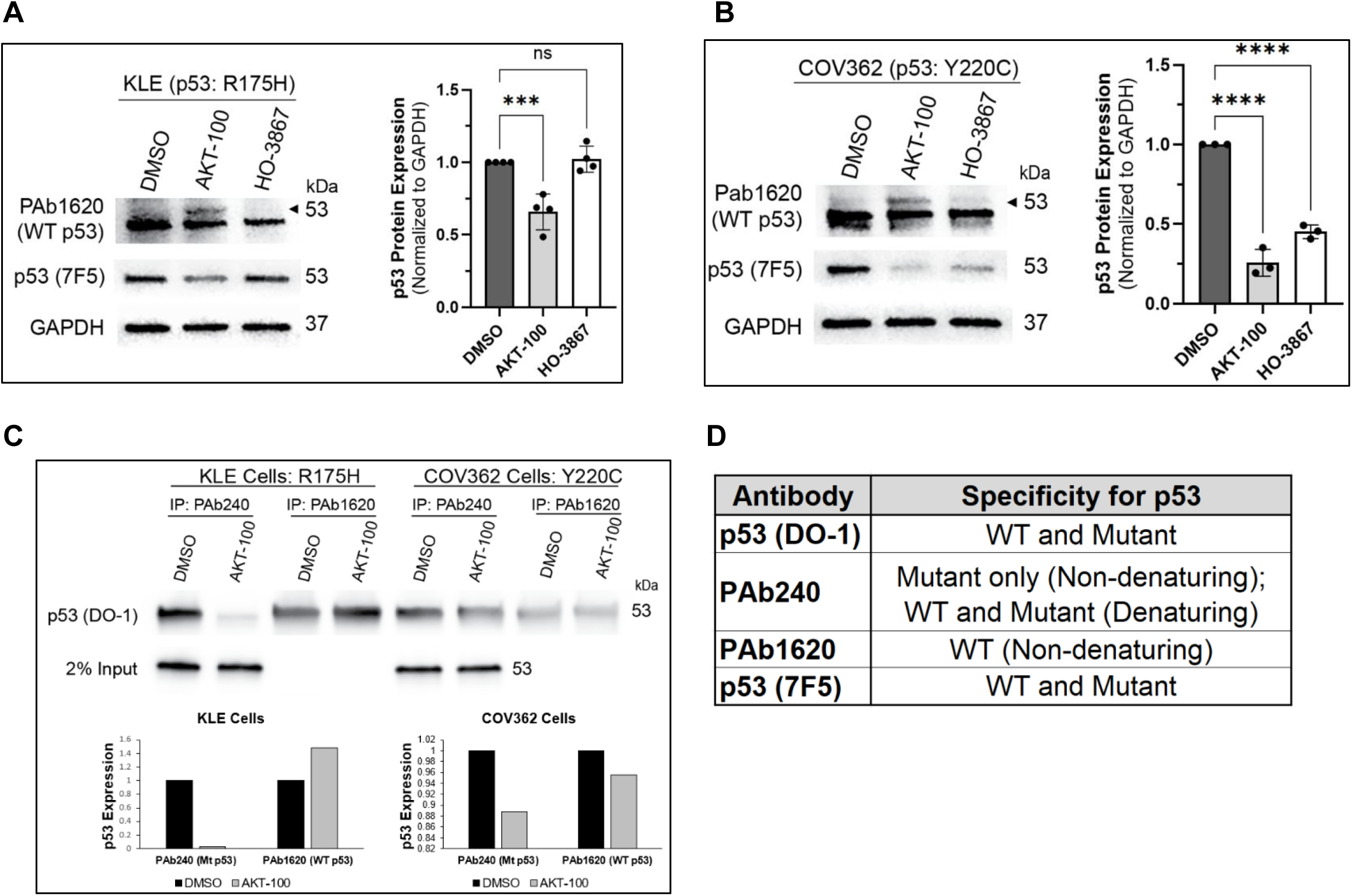
Mutant p53 protein is degraded while WT p53 is reactivated upon treatment with AKT-100. **(A)** Western blot analysis showing reactivation of PAb1620 WT p53 (band with arrow) while subsequently decreasing mutant p53 in KLE cells treated with AKT-100 for 24 hours. Note: Lower bands in PAb1620 are non-specific under denaturing conditions. Quantification (n=4) is shown on the right for degradation of mutant p53 (p53 (7F5) antibody). **(B)** Western blot showing degradation of mutant p53 in COV362 cells upon treatment with both AKT-100 and HO-3867. Quantification (n=3) is shown on the right. **(C)** Immunoprecipitation comparing mutant p53 (PAb240) versus WT p53 (PAb1620) demonstrates reactivation of WT p53 after treatment with AKT-100 for 24 hours in KLE and COV362 cells. **(D)** Table listing p53 antibody specificity under denaturing and non-denaturing conditions used in experiments. One-way ANOVA was performed to compare to DMSO treatment, where statistical significance is indicated as follows: ***, p < 0.001; ****, p<0.0001; ns, not significant.

The induction of pro-apoptotic genes is a consistent effect of curcumin analogues including AKT-100. As displayed in Figure 3B (right panel), endometrial cell lines KLE and AN3CA had an increase in apoptosis genes *BBC3* (PUMA) and *PMAIP1* (NOXA), whereas all cell lines tested increased multiple ER stress genes (*ERN1*, *XBP1*, *EIF2AK3*, *HSPA5*, *PPP1R15A*, *ATF4* and *CHOP*). Notably, prolonged ER stress activates apoptotic pathways contributing to the goal of inhibiting cancer cell proliferation (55). To understand more of the impacts of AKT-100 on the many other cancer-related pathways, we report additional anti-cancer transcriptomic effects in Supplemental Figure 4.

In addition, since it was shown that AKT-100 also binds STAT3 (Figure 1F) but has little effect on STAT3 expression except in AN3CA cells (Figure 3B, right panel), we asked whether AKT-100 inhibits STAT3 activation by altering phosphorylation. Indeed, Western blot analysis confirms that AKT-100 functions to also inhibit STAT3 activation due to a decrease in phosphorylated STAT3 at Tyrosine 705 in both KLE and COV362 cells (Figure 3D). The two major mechanisms of STAT3 activation are through phosphorylation at Tyrosine 705 which drives nuclear activation for DNA binding that leads to transcription of target genes or mitochondrial activation via Serine 727 phosphorylation (7).

We find that AKT-100 reactivates WT p53 expression based on immunoprecipitation and Western blot analysis using mutant and WT-specific antibodies. Figure 4A-C demonstrate that AKT-100 increased WT p53 expression in both KLE and COV362 cells. According to the literature (56–58), PAb1620 is specific for recognizing WT p53, and PAb240 will only recognize mutant p53 under non-denaturing conditions. In these experiments, Figure 4A shows a reduction in mutant p53 only with AKT-100 in KLE cells based on an antibody that recognizes both WT and mutant p53. However, mutant p53 is degraded upon treatment with both AKT-100 and HO-3867 in COV362 cells (Figure 4B). Immunoprecipitation with mutant-specific antibody PAb240 in KLE and COV362 cells show capture of only mutant p53 in DMSO conditions, while pull-down with WT-specific PAb1620 shows more isolation of WT p53 with AKT-100 treatment. These results demonstrate that AKT-100 functions as a WT p53 reactivator.

### AKT-100 is synergistic with both a PARP inhibitor and with chemotherapy

As single agents, AKT-100 and HO-3867 are effective at low µM concentrations in reducing cancer cell proliferation (Figure 2). On the other hand, we determined that the PARP inhibitor olaparib had high IC_50_ values of 10.48 µM in OVCAR3 and 65.11 µM in COV362 cells (Figures 5A, 5B, 5I), signaling PARP inhibitor resistance. Thus, we asked whether we could improve this treatment by combining it with the AKT-100 curcumin analogue. Indeed, as shown in Figure 5A, addition of 400 nM AKT-100 reduced the IC_50_ value of olaparib in combination to 4.73 µM (Figure 5I) when combined with olaparib. Also shown in Figure 5B is that addition of 20 µM constant olaparib to increasing concentrations of AKT-100 reduced the IC_50_ value of AKT-100 to 0.07 µM when used in combination. The combinatorial index was calculated to be 0.75 as shown in Figure 5I, and values less than 1 indicate synergism was achieved with combination drugging. The experiment was repeated in COV362 cells, and compared to olaparib alone, the combination with 150 nM AKT-100 significantly reduced the IC_50_ value of olaparib from 65.11 to 15.31 µM (Figure 5C). The reverse combination, with 30 µM constant olaparib added to increasing concentrations of AKT-100 reduced the AKT-100 IC_50_ value from 0.20 µM to 0.07 µM, and again the combinatorial index was 0.59, indicating synergy. Overall, these data demonstrate that the combination of olaparib and AKT-100 significantly improves cell killing compared to either agent alone, and it is possible to reduce the drug concentrations significantly and still achieve therapeutic responses. Considering that olaparib is most effective in cancers with hereditary BRCA1/BRCA2 mutations as well as in cases of tumor somatic mutations in homologous recombination repair genes, the inhibitory impact of AKT-100 on genes involved in homologous recombination shown in Figure 4D likely underlies the synergy when combined with olaparib, basically creating cancer cell homologous recombination repair deficiency.

**Figure 5.**
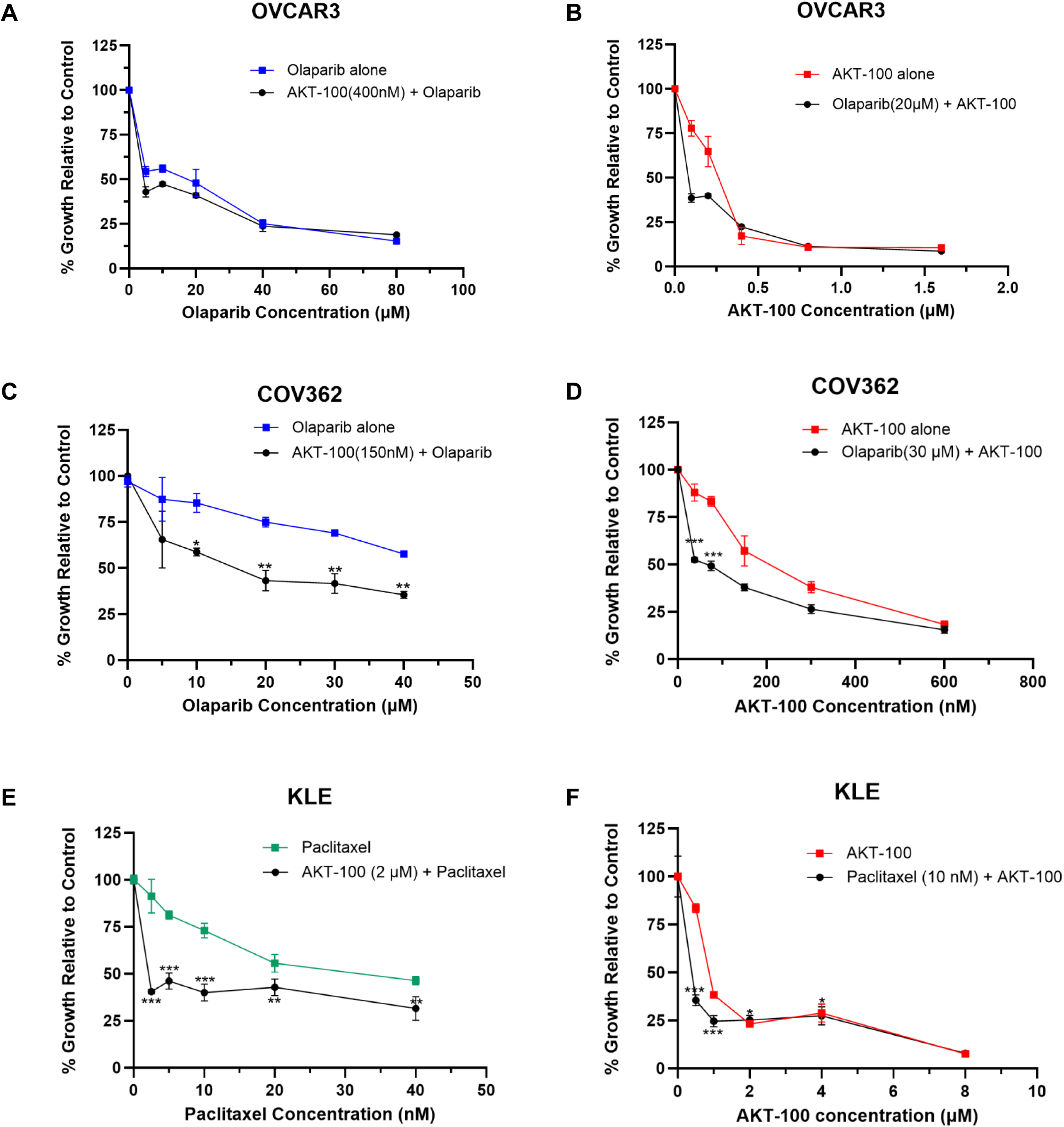
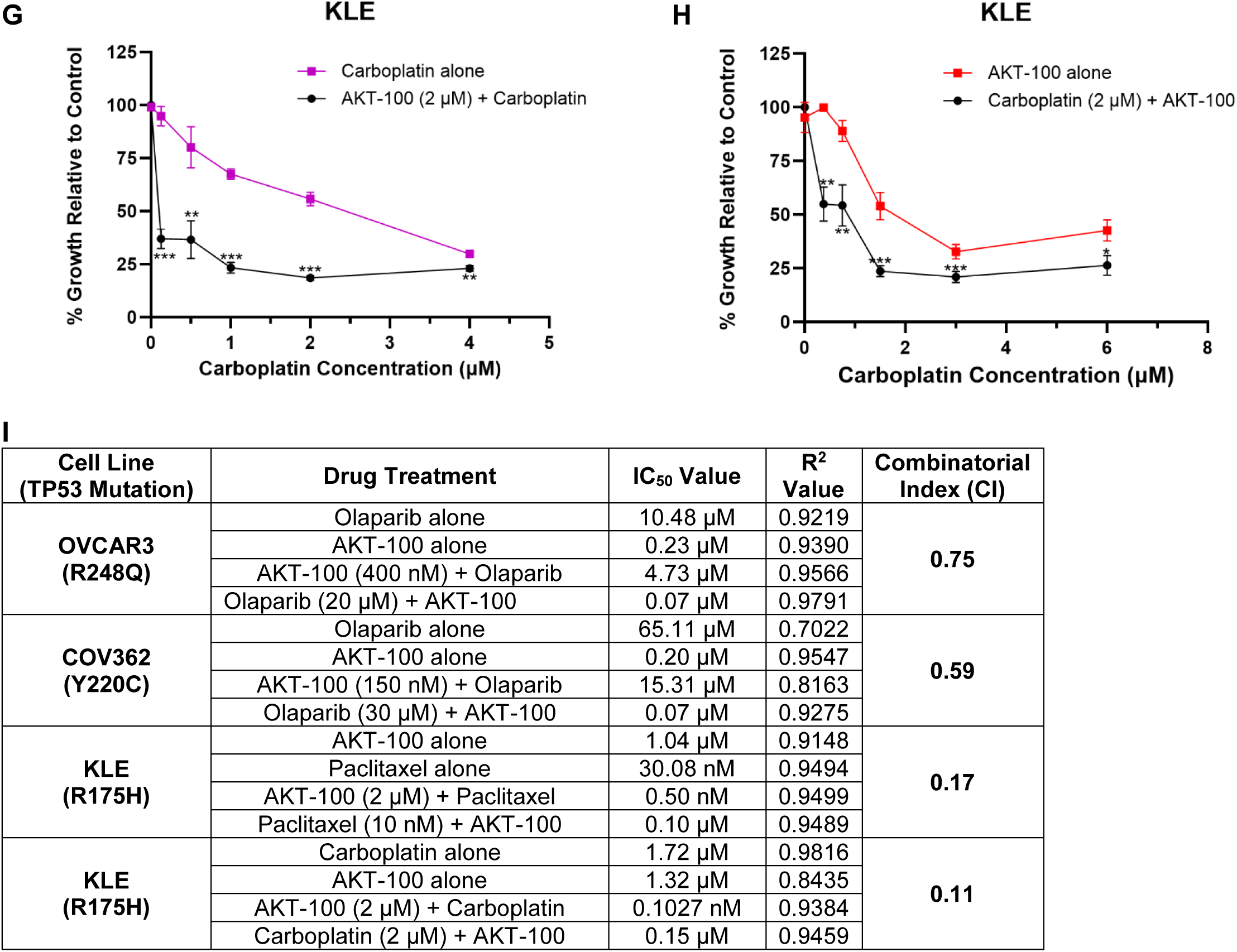
AKT-100 is synergistic with both a PARP inhibitor and with chemotherapy. **(A)** Growth inhibition curves in OVCAR3 cells comparing olaparib alone versus olaparib with 400 nM AKT-100. **(B)** Cell proliferation assay displaying OVCAR3 cells drugged with AKT-100 alone compared to AKT-100 plus constant 20 µM olaparib. **(C)** Cell proliferation assay displaying COV362 cells drugged with olaparib alone versus olaparib with 150 nM AKT-100. **(D)** Proliferation assay comparing AKT-100 alone versus AKT-100 plus 30 µM olaparib in COV362 cells. **(E)** KLE cells were treated with paclitaxel alone versus paclitaxel plus 2 µM constant AKT-100 or **(F)** AKT-100 alone compared to AKT-100 with 10 nM paclitaxel. **(G)** Proliferation assay in KLE cells treated with carboplatin alone or carboplatin with 2 µM constant AKT-100. **(H)** Graph comparing cell proliferation of KLE cells treated with either AKT-100 alone versus AKT-100 with 2 µM carboplatin. **(I)** Table listing the combination drug treatments in OVCAR3, COV362, and KLE cells treated with chemotherapy drugs and AKT-100. Listed are determined IC_50_ values, R^2^ values and calculated combinatorial indices. For A-H, single drug treatments are blue lines and combination treatments are red lines. Unpaired t-tests were performed to compare the two groups, and statistical significance is indicated as follows: *, p < 0.05; **, p < 0.01; ***, p < 0.001.

We next asked whether other chemotherapeutic regiments such as paclitaxel and carboplatin, that induce DNA damage, are also synergistic with AKT-100. We tested KLE cells with paclitaxel and determined the IC_50_ to be 30.08 nM, but addition of 2 µM AKT-100 reduced this value to 0.50 nM (Figures 5E, 5I). The reverse combination showed the IC_50_ for AKT-100 alone was 1.04 µM, however addition of only 10 nM of constant paclitaxel dropped the IC_50_ of AKT-100 10-fold to 0.10 µM (Figures 5F, 5I). The combinatorial index was calculated to be 0.11─well below 1 (Figure 5I). As for combination drugging with carboplatin, we again determined it to be synergistic in KLE cells which harbor the most resistant p53 R175H mutation. Treatment with carboplatin alone demonstrated an IC_50_ of 1.72 µM, and addition of 2 µM AKT-100 reduced this to 0.1027 nM (Figures 5G, 5I). The reverse combination determined an IC_50_ with AKT-100 alone to be 1.32 µM and combination with 2 µM carboplatin reduced it to 0.15 µM (Figure 5H, 5I). The combinatorial index was calculated to be 0.11 (Figure 5I), demonstrating significant synergy between AKT-100 and carboplatin.

## Discussion

Despite the overall improvement in outcomes for many cancer patients today, women suffer from increased year over year incidences (59) and deaths (60) from uterine endometrial cancer up to the current decade. Ovarian cancer, now understood to arise primarily in the epithelium of the fallopian tube and to spread early in the process of oncogenesis, is often lethal with fewer than 50% of patients surviving 5 years after diagnosis (61). A common thread among these malignancies is the inactivation of the central tumor suppressor p53, mostly through mutation. We and others have evaluated the rate of the various types of mutations in the p53 gene, *TP53*, and found that approximately 70 percent result in single amino acid hotspot alterations in the DNA binding domain of p53, preventing its normal pro-apoptotic and DNA repair transcriptional activity, while enhancing some mutant p53 functions that encourage oncogenesis and tumor progression (62). Recurrent mutations found in nearly all types of cancers include R175H, R248Q/W, Y220C, R273H/C, R282W, G245S, and R249S. Many of the recurrent missense mutations occur at methylated CpG sites which encode arginine residues that contact the DNA and are conserved over evolutionary scales (4).

Indeed, considering comparative biology and evolution, the critical importance of functional, WT p53 in cancer prevention and longevity is emphasized when studying other animals that are resistant to cancer and live longer than expected lives (2). These include elephants and bats, with genomes encoding multiple p53 genes – 20 copies of *TP53* are encoded in the elephant genome, and two copies of *TP53* exist in the bat genome compared to a single copy in most species including humans. The frequent mutation of *TP53* creates individual and species vulnerabilities to cancer; however, could there be a therapeutic mechanism to reverse or prevent such mutations and to enhance a longer and healthier lifespan? The goal of our research is to develop such a strategy.

The molecular properties of curcumin as an anti-inflammatory, antioxidant, anti-angiogenic and anti-cancer agent have been the subject of investigations over decades, as summarized by Vander Jagt *et al*. (29). One of the most important observations has been the identification of p53 as a target for the curcumin analogue HO-3867 (26, 44). This supports the likelihood that curcumin and its analogues may serve as p53 reactivators with the potential to bind to mutant forms of p53 to recapitulate wild type pro-apoptotic activity in cancer cells. Several p53 reactivator therapeutic molecules have been extensively assessed in preclinical studies, such as HO-3867, and two therapeutics, APR-246 and rezatapopt (PC14586), have been tested in clinical trials (63, 64). Rezatapopt, which is specifically designed to inhibit a single mutant, p53 Y220C, has shown activity in a phase I trial of patients whose tumors expressed that specific mutation, including patients with endometrial cancer who responded (65). The advantage of a curcumin analogue is the broader range of p53 mutant targeting beyond a single mutation; hence, the potent anticancer activity of such agents continues to bode well for the eventual development and use of curcumin analogues in medicine.

In these studies, we report the activity of AKT-100, a novel derivative of HO-3867 and a curcumin analogue, as an anticancer therapeutic with single agent activity. We demonstrate the substantial impact of relatively low concentrations of AKT-100 on cell viability using multiple cell models of advanced gynecologic cancer with mutant forms of p53. Our data confirm that mutant p53 proteins are bound by AKT-100, and that this results in the re-establishment of WT p53 transcriptional effects including the induction of apoptotic genes, the enforcement of cell cycle checkpoints and the inhibition of multiple cancer cell DNA repair pathways. We have also shown that AKT-100 inhibits the phosphorylation of STAT3 at Tyrosine 705, thus further preventing oncogenic activity while WT p53 is reactivated. A single agent that can simultaneously avert oncogenesis by stopping STAT3 activity while re-establishing tumor suppressive functionality of WT p53 could be a major advance in cancer therapy.

Additionally, as a result of the inhibition of DNA repair pathways, AKT-100 is synergistic with the PARP inhibitor olaparib and with paclitaxel and carboplatin, the standard chemotherapeutic agents used to treat p53 mutant and other advanced forms of endometrial and ovarian cancers. The opportunity to prevent resistance to chemotherapy and to PARP inhibitors using a curcumin analogue is highly clinically relevant (66). Additionally, a strength of the work reported herein is the identification of a novel curcumin analogue that binds to mutant forms of p53 with the potential to reactivate the WT p53 transcriptome. We and others have proposed that reactivating the normal, pro-apoptotic effects of WT p53 in tumor cells harboring mutant forms of p53 can have substantial clinical benefit for patients with advanced cancers. Nevertheless, given the reported therapeutic impact of curcumin analogues in diseases ranging from cancer to arthritis to Alzheimer’s disease, multiple mechanisms of action are highly likely and should be delineated in follow-up studies (33). Other targets, including STAT3, as reported to be an additional target of HO-3867 beyond p53 (67), are likely to be involved in the activity of AKT-100 and other curcumin analogues, and these targets should be more fully defined. In addition, identifying curcumin analogues in future studies with a higher affinity for multiple mutant forms of p53 compared to WT would be advantageous (20). With the accomplishment of these goals, the future may hold the promise of synthesizing different curcumin analogues with activities tailored to inhibit targets promoting cancer as well as other diseases.

## Acknowledgements

This work was completed with support from the UNMCCC Biostatistics, ATG Genomics and the UNM Flow Cytometry Core Shared Resources. Graphical abstract was created with BioRender.com.

## Authorship contribution

**Geneva Williams:** Writing – original draft, Methodology, Investigation, Formal analysis, Data curation. **Lane E. Smith, Jamie L. Padilla, Alexander Goss, Daisy Belmares-Ortega, Xiangxiang Wu, Jennifer N. Daw, Prakash Jagtap, Josh K. Monts:** Methodology, Investigation, Data curation. **Jun-yong Choe:** Methodology, Investigation, Formal analysis, Manuscript development. **Sumegha Mitra**: Writing - review and editing of the manuscript. **Kimberly K. Leslie:** Writing – review & editing, Writing – original draft, Validation, Supervision, Resources, Project administration, Methodology, Investigation, Funding acquisition, Formal analysis, Data curation, Conceptualization.

## Data availability statement

The data supporting the findings of this study are available within the article and its supplementary materials. The RNA sequencing data performed in this study are uploaded and available through GEO, accession number GSE310454.

