## Supplemental Information for "The novel curcumin analogue AKT-100 targets both mutant p53 and STAT3 in gynecologic cancer cells"

### Supplemental Figure 1

***Synthetic pathways for the preparation of AKT-100, AKT-111, AKT-121, AKT-109 and AKT-115.***

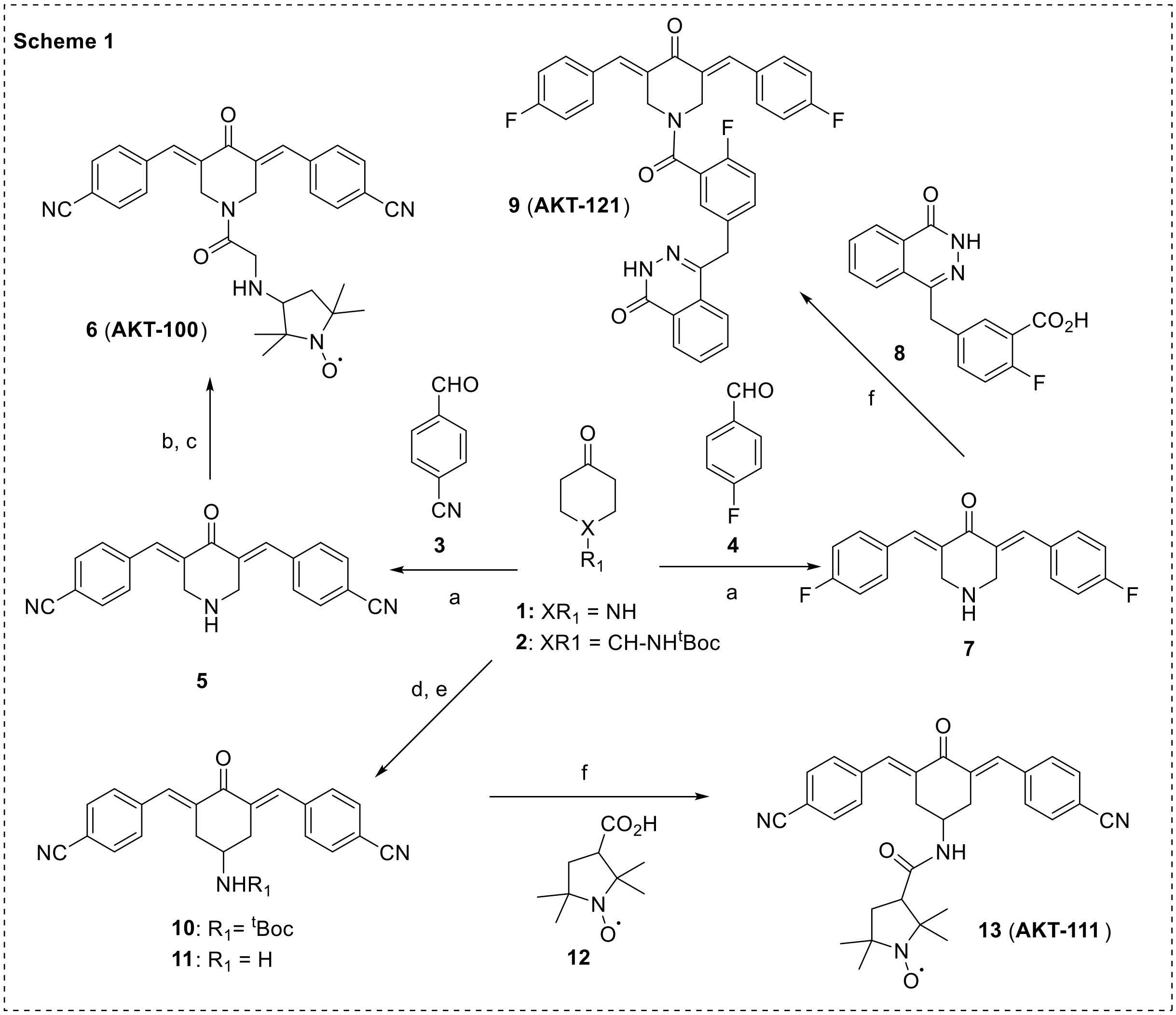


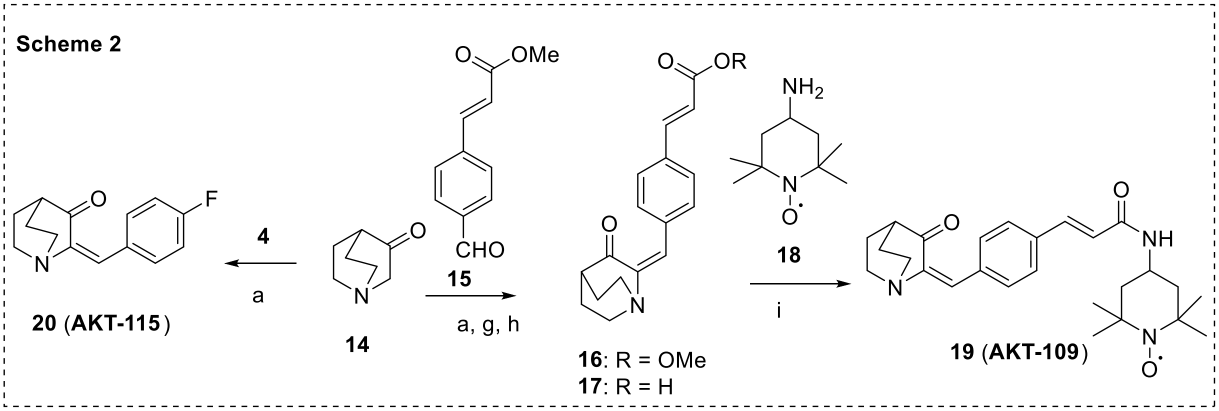


**Reagents and reaction conditions**: (a) $BF_3$-$Et_2O$, 0 $^oC$, 8 hr; (b) $ClCOCH_2Cl$, $NE_3$, $CH_2Cl_2$, rt, 5 hr; (c) 3-aminoproxyl, DMF, DMAP, $NEt_3$; (d) NaOH, EtOH, rt, 3 hr; (e) 4N HCl, MeOH, THF, rt, 6 hr; (f) $T_3P$ reagent (propane phosphoric acid anhydride, DMF, DIPEA; (g) LiOH, MeOH, THF; (h) 2N HCl; and (i) DMF, HATU, DIPEA, 2 hr, rt

### Materials and methods:

Proton nuclear magnetic resonance ($^1$H-NMR) spectra were obtained from Varian 300 MHz spectrophotometer. Thin layer chromatography, TLC, was carried out on precoated TLC plates with silica gel 60 F-254 and preparative TLC on precoated Whatman 60A TLC plates. All intermediates were characterized on the basis of NMR ($^1H$). $^1H$-NMR of the final paramagnetic nitroxide compounds are significantly shifted and broadened spectral lines due to the unpaired electron's magnetic moment. These compounds were analyzed by Mass spectral (MS) analysis data. All starting materials (compound # 1, 2, 3, 4, 12, 14, 15 and 18), solvents and reagents were purchased from Sigma-Aldrich, USA. 2-Fluoro-5-((4-oxo-3,4-dihydrophthalazin-1-yl)methyl)benzoic acid (compound # 8, CAS No.: 763114-26-7) was purchased from Ambeed (Buffalo Grove, IL 60089, USA). Flash column chromatography was performed with Merck ultra-pure silica gel (Darmstadt, Germany, 230–400 mesh). Analytical HPLC was performed using a Waters Alliance 2795 series system, equipped with a Waters UV 2996PAD (set at 254 nM) and Micromass MS Quattro LC detector or using Waters Alliance 2690 series system, equipped with the Micromass LCT detector. A YMC-Pack-ODS-AQ (series AQ12505-1546WT, size 150 mm X 4.6 mm, S-5 µM) column was used. Typically, a gradient mobile phase starting with 70% water with 0.05% ammonium formate and 30% methanol with 0.05% ammonium formate. Flow rate 0.8 ml/min. Based on the UV and MS detectors, all final compounds which evaluated in the in vitro assays showed HPLC purity >95%.

**Compound 5**: 4-Piperidone monohydrate hydrochloride 1 (1 mol eq.) and 4-formyl benzonitrile **3** (2.2 eq) and acetonitrile (ACN) were added into reactor at room temperature. BF3.Et2O (4 mol. eq) was added drop-wise at 20-30°C in one portion. The reaction mixture was refluxed for 8 h at 70°C. Reaction mixture was cooled to 10-20 °C and stirred for 1-2 hours. The solid was filtered and washed with ACN to afford aldol product compound **5**. It was dried under vacuum at 40-50° and was isolated in 80 % yield.

**Compound 6** (**AKT-100**): Compound **5** (1 mol. eq) was suspended in ethyl acetate. Saturated NaHCO3 solution was added, followed by chloroacetyl chloride (5 eq). The reaction mixture was stirred vigorously for 2 hours and the resulting solid collected by filtration, washed with water and ethyl acetate. The solid was dissolved in dichloromethane and DMF (15 ml each) and 1.1 eq of 3-aminoproxyl was added. Upon stirring 2h at 50°C and 4 hr at ambient temperature the reaction mixture was concentrated, and the residue purified by column chromatography using 5-20% MeOH-$CH_2Cl_2$ to give Compound **6** (AKT-100) in 52% yield. LCMS: 523.2 [M+H]+.

**Compound 10**: tert-Butyl (4-oxocyclohexyl) carbamate **2** (1.07 g, 5.0 mmol) and 4-formyl benzonitrile (1.4 g, 2.0 eq) and EtOH (10 ml) were added into reactor at room temperature. NaOH (200 mg, 5.0 mmol) was added in one portion. The reaction mixture was stirred for 3 hr at room temperature. The yellow solid was filtered and washed with water and EtOH to give compound **10** (1.5 g, 3.42 mmol). LCMS: 440.2 $[M+H]^+$.

**Compound 11**: Compound **10** (610 mg, 1.39 mmol) was dissolved in MeOH/THF (20 ml/20ml) and cooled to 0°C. 4N HCl (10 ml) in dioxane was added in one portion. The reaction mixture was stirred at ambient temperature for 6 h. The resulting solid collected by filtration, washed with ethyl acetate to give compound **11** (460 mg, 1.23 mmol). LCMS: 340.2 $[M+H]^+$.

**Compound 13** (**AKT-111**): Compound **11** (397 mg, 1.06 mmol) and compound **12** (218 mg, 1.06 mmol) were suspended in DMF (5 ml). $T_3P$ reagent (1.2 ml g, 1.0 eq) was added, followed by diisopropylethylamine (DIPEA) (0.82 ml, 3 eq). The reaction was stirred overnight at room temperature. The reaction mixture was poured into water. The resulting solid was collected by filtration, washed with water and acetonitrile. The desired compound 11 (AKT-111) was obtained (350 mg, 0.69 mmol). HPLC purity: > 98%. LCMS: 508.2 $[M+H]^+$.

**Compound 7**: 4-Piperidone monohydrate hydrochloride **1** (1.53 g, 10 mmol eq.) was placed in a 100 ml round bottom flask and cooled to 0°C. $BF_3.Et_2O$ (10 ml) was added dropwise followed by 4-fluorobenzaldehyde **4** (2.1 ml, 2.0 eq) in one portion. The reaction mixture was stirred for overnight at room temperature. The reaction was carefully quenched with a saturated solution of NaHCO3. The light-yellow solid was filtered and washed with water and EtOH to give intermediate compound **7**. LCMS: 312.2 $[M+H]^+$.

**Compound 9** (**AKT-121**): The compound **7** (311 mg, 1.0 mmol) and 2-fluoro-5-((4-oxo-3, 4-dihydrophthalazin-1-yl) methyl) benzoic acid **8** (298 mg, 1.0 mmol) were suspended in DMF (5 ml). $T_3P$ (propane phosphonic acid anhydride) reagent (1.0 ml g, 1.0 eq) was added, followed by diisopropylethylamine (DIPEA) (0.52 ml, 3 eq). The reaction was stirred at room temperature. After 1h, the reaction mixture became a clear solution. The mixture was poured into water. The resulting solid was collected by filtration, washed with water, and methanol. The desired compound **9** was obtained (300 mg). HPLC purity: > 98.5%. LCMS: 592.2 $[M+H]^+$.

**Compound 16**: Quinuclidin-3-one hydrochloride **14** (753 mg, 4.66 mmol eq.) was placed in a 100 ml round bottom flask and cooled to 0°C. $BF_3.Et_2O$ (10 ml) was added dropwise followed by methyl (E)-3-(4-formylphenyl) acrylate **15** (887 mg, 1.0 eq) in one portion. The reaction mixture was stirred for 8 hrs at room temperature. The reaction was carefully quenched with a saturated solution of NaHCO3. The light-yellow solid was filtered and washed with water and EtOH to give compound **16** (700 mg 2.36 mmol). LCMS: 298.2 $[M+H]^+$.

**Compound 17**: Methyl (E)-3-(4-((Z)-(3-oxoquinuclidin-2-ylidene) methyl) phenyl) acrylate **16** (700 mg, 2.35 mmol) was dissolved in THF (30 ml) and MeOH (10 ml) at room temperature. LiOH (1N 10 ml) was added and stirred for overnight at room temperature. THF and MeOH were evaporated under reduced pressure. The residue was acidified with HCl (2N) to PH around 3. The yellow solid was filtered and washed with water to give compound **17** (550 mg, 1.94 mmol). LCMS: 284.2 $[M+H]^+$.

**Compound 19** (**AKT-109**): Compound **15** (545 mg, 1.93 mmol) and compound **18** (330 mg, 1.93 mmol) were suspended in DMF (5 ml). HATU reagent (740 g, 1.0 eq) was added, followed by diisopropylethylamine (DIPEA) (1.0 ml, 3 eq). The reaction was stirred 2 hrs at room temperature. The reaction mixture was concentrated and purified with reverse-phase HPLC to give compound **19** (AKT-109) (400 mg, 0.92 mmol). HPLC purity: > 98%. LCMS: 437.4 $[M+H]^+$.

**Compound 20** (**AKT-115**): Quinuclidin-3-one hydrochloride **14** (753 mg, 4.66 mmol eq.) was placed in a 100 ml round bottom flask and cooled to 0°C. BF3.Et2O (10 ml) was added dropwise followed by 4-fluoro-benzaldehde **4** (1.0 eq) in one portion. The reaction mixture was stirred for 8 hrs at room temperature. The reaction was carefully quenched with a saturated solution of NaHCO3. The light-yellow solid was filtered and washed with water and EtOH to give compound **20** (750 mg). LCMS: 232.1 [M+H]+.

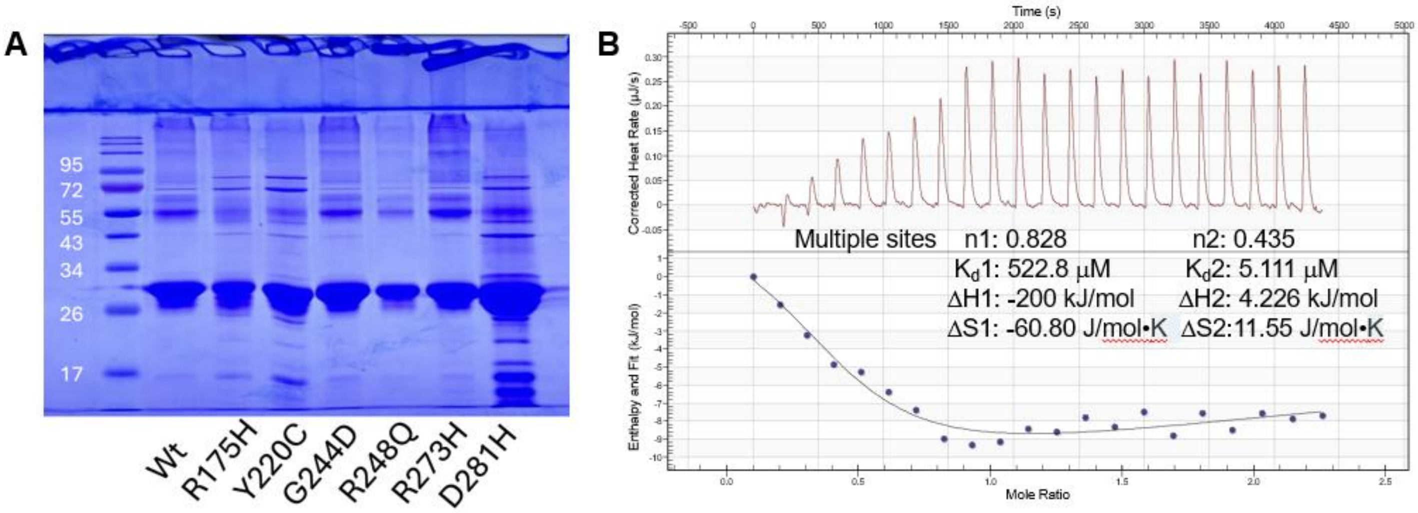


**Supplemental Figure 2. AKT-100 binds wild type and mutant p53 proteins. (A)** SDS PAGE gel of purified p53 wild-type and indicated mutants. Loaded protein samples are the result of a single step purification on a Talon column. **(B)** ITC thermogram of p53 upon AKT-100 titration. The ITC predicts a $K_d$~ 5.1 and 523 μM for AKT-100 binding to p53. Triplicate ITC measurements all produced the same profile as shown.

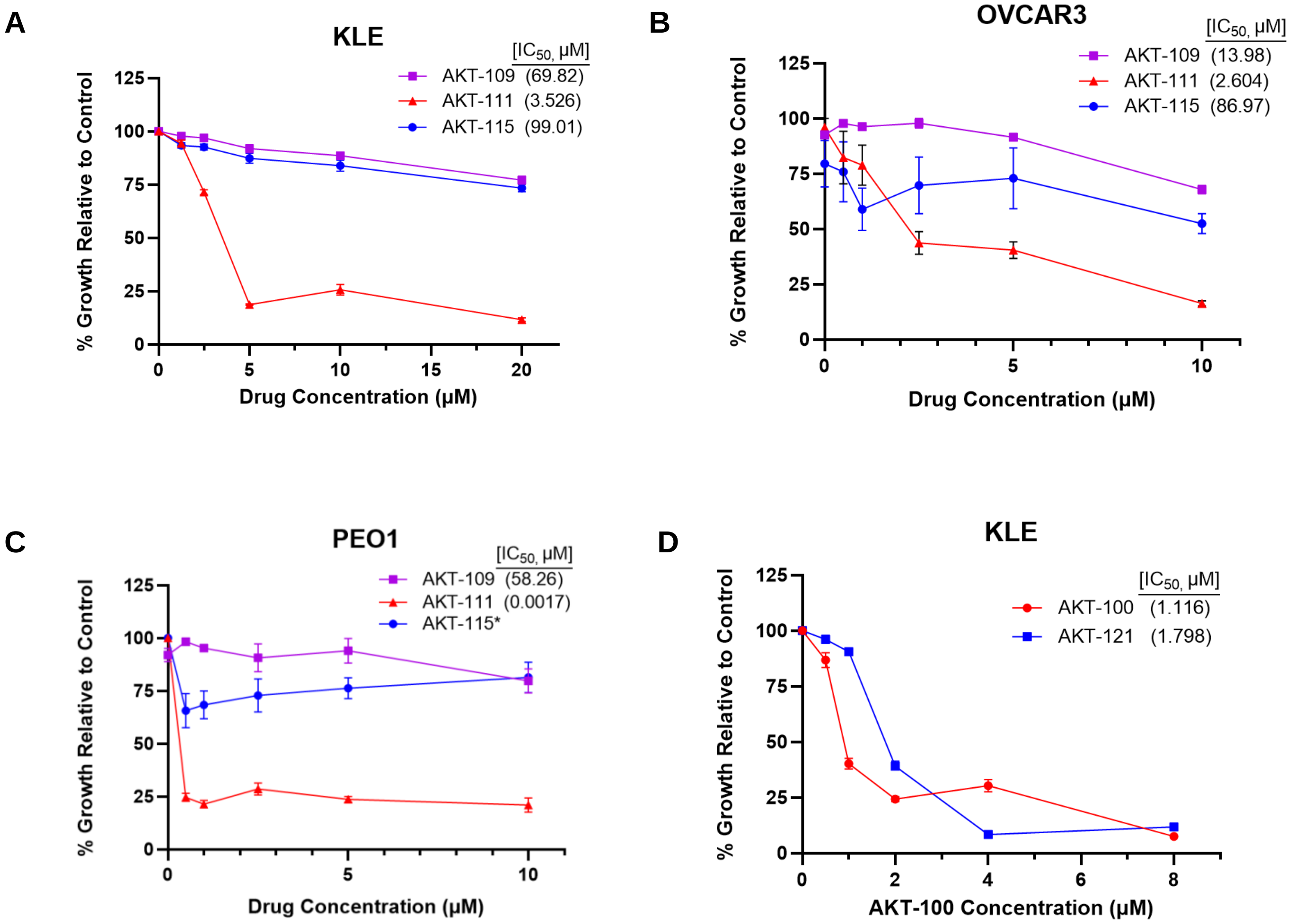

**E**

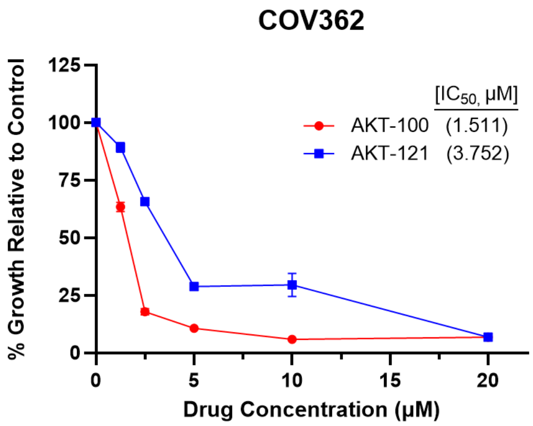


**Supplemental Figure 3. AKT-111 and AKT-121 also demonstrate potential activity against cancer cells.** Cell proliferation assays for **(A)** KLE **(B)** OVCAR3 and **(C)** PEO1 cells drugged with AKT derivatives AKT-109, AKT-111 and AKT-115. *Note: The IC50 for AKT-115 was unable to be obtained. Cell proliferation assays for **(D)** KLE and **(E)** COV362 cells drugged with AKT-100 versus AKT-121. Error bars depict standard error of the mean for three replicate wells.

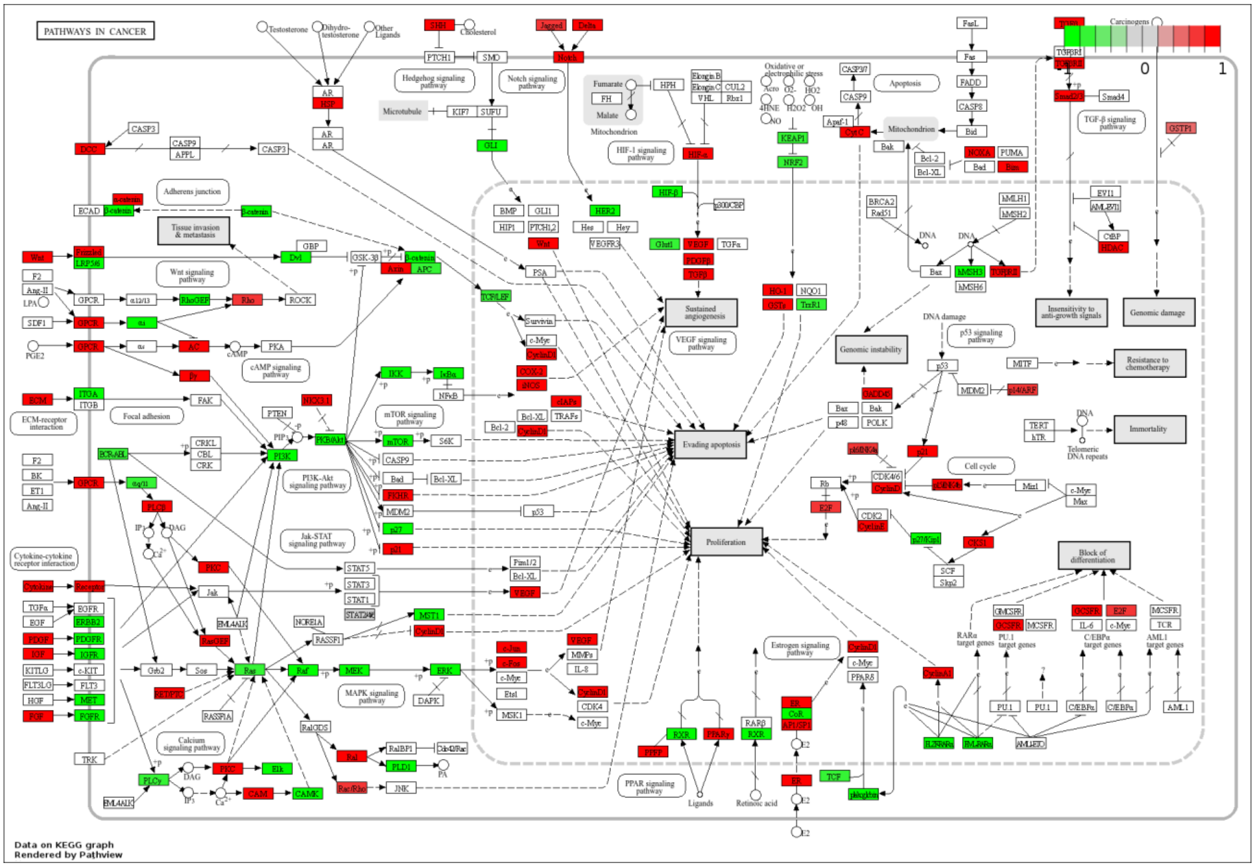


**Supplemental Figure 4. Pathview analysis of RNA sequencing results in KLE cells drugged with AKT-100 showing pathways in cancer.** Upregulated genes are highlighted in red and downregulated genes are highlighted in green.

| RNA Sequencing (24 Hours) | | KLE (R175H) | | COV362 (Y220C) | | OVCAR3 (R248Q) | |
|---|---|---|---|---|---|---|---|
| | | AKT-100 | HO-3867 | AKT-100 | HO-3867 | AKT-100 | HO-3867 |
| Protein Name | Gene | Log$_2$ Fold Change | | | | | |
| p21 | *CDKN1A* | 2.64 | 2.14 | 3.31 | 3.51 | 1.40 | |
| GADD45 | *GADD45A* | | 3.81 | 1.34 | 2.11 | 1.75 | |
| | *GADD45B* | 5.13 | 2.72 | 4.34 | 2.93 | 3.60 | |
| | *GADD45G* | 12.95 | | 6.02 | 3.34 | 3.72 | |

**Supplemental Figure 5. Comparison of cell cycle regulating transcripts in cells treated with AKT-100 versus HO-3867.** Chart comparing log$_2$ fold transcript change values of cell cycle regulators p21 and GADD45 in KLE, COV362 and OVCAR3 cells treated with AKT-100 versus HO-3867.
